# An ALK1-governed monocytic lineage shapes an immunosuppressive landscape in breast cancer metastases

**DOI:** 10.1101/2024.06.15.599147

**Authors:** Mehrnaz Safaee Talkhoncheh, Jonas Sjölund, Paulina Bolivar, Ewa Kurzejamska, Eugenia Cordero, Teia Vallès Pagès, Sara Larsson, Sophie Lehn, Gustav Frimannsson, Viktor Ingesson, Sebastian Braun, Jessica Pantaleo, Clara Oudenaarden, Martin Lauss, R. Scott Pearsall, Göran B. Jönsson, Charlotte Rolny, Matteo Bocci, Kristian Pietras

## Abstract

The biology centered around the TGF-β type I receptor ALK1 (encoded by *ACVRL1)* has been almost exclusively based on its reported endothelial expression pattern since its first functional characterization more than two decades ago. Here, in efforts to better define the therapeutic context in which to use ALK1 inhibitors, we uncover a population of tumor-associated macrophages (TAMs) that, by virtue of their unanticipated *Acvrl1* expression, are effector targets for adjuvant anti-angiogenic immunotherapy in mouse models of metastatic breast cancer. The combinatorial benefit depended on ALK1-mediated modulation of the differentiation potential of bone marrow-derived granulocyte-macrophage progenitors, the release of CD14^+^ monocytes into circulation, and their eventual extravasation. Notably, *ACVRL1*^+^ TAMs coincided with an immunosuppressive phenotype, and were over-represented in human cancers progressing on therapy. Accordingly, breast cancer patients with a prominent *ACVRL1*^hi^ TAM signature exhibited a significantly shorter survival. In conclusion, we shed light on an unexpected multimodal regulation of tumorigenic phenotypes by ALK1 and demonstrate its utility as a target for anti-angiogenic immunotherapy.

**Graphical abstract:** See submitted file

## Introduction

The activity of the transforming growth factor (TGF)-β superfamily of ligands and receptors constitutes an essential component of physiological and developmental processes (1). In a similar fashion —owing to the pleiotropic effects of this complex pathway— the molecular cues instigated by TGF-β signaling are an established hallmark of cancer (2). Nonetheless, translation to clinical application of TGF-β pathway inhibitors has proved difficult, since the signal transduction cascade is cell type-specific (3), environmentally controlled (4), and stage-dictated throughout tumorigenesis (5). Knock-out studies in mice have detailed the normal function of the TGF-β type I receptor Activin Receptor-Like Kinase (ALK)1, invariably resulting in embryonic lethality due to gross vascular alterations (6). Even though additional studies reported antithetic roles for ALK1 during physiological angiogenesis, a more generalized consensus has been attained in cancer, where ALK1 mediates pro-angiogenic stimuli linked to disease evolution (7, 8).

We and others have previously exploited targeting of ALK1 activity to impinge on tumor growth, either through genetic or pharmacological means (9–11), *e.g. via* the ligand trap RAP-041/dalantercept (referred to as ALK1-Fc from hereon), which binds the circulating high-affinity ligands bone morphogenetic protein (BMP)-9 and -10 (9, 12–14). Upon ALK1 blockade, tumor-associated vasculature is pruned, display an increased pericyte coverage (conceivably contributing to a more efficient drug delivery), with a resulting reduction of metastatic dissemination to the lungs (9, 13). Despite these promising results, the clinical endorsement of ALK1 inhibitors was short-lived: indeed, following promising phase 1 trials in patients with heavily pre-treated advanced solid cancers (15, 16), expansion and phase 2 tests with dalantercept, either alone or in combination with Vascular Endothelial Growth Factor (VEGF) inhibitors, fell short of additional favorable effects, even in presence of a tolerable safety profile (17–22). More recently, we determined that the network of genes correlated with *ACVRL1* expression is associated with a series of processes defining the tumor microenvironment (TME) across human solid malignancies, with a large overrepresentation of pathways linked to immune cell function and regulation (23). However, how ALK1 signaling in the tumor endothelium and beyond contributes to metastatic and drug-resistant properties of the local tumor *milieu* remains unexplored.

Here, by modeling adjuvant (*i.e.* post-surgical) therapy targeting ALK1 in experimental metastatic breast cancer, we characterize a previously overlooked subpopulation of tumor-associated macrophages (TAMs) identified by expression of *Acvrl1* (encoding for ALK1). Through integration of *in vivo* platforms, *ex vivo* assays and longitudinal *in silico* data, we demonstrate that *Acvrl1*^+^ TAMs derived from circulating CD14^+^ monocytes that were recruited to the tumor site. Indeed, blunted ALK1 signaling directly shaped the potency of hematopoietic progenitor cells (HPCs) in the bone marrow (BM) by hindering their differentiation into the monocytic lineage, as well as the subsequent mobilization of monocytes and their maturation to macrophages. Moreover, the emergence of *ACVRL1*^+^ TAMs was associated with disease progression upon development of therapeutic resistance in cancer patients. Consequently, high expression of a TAM-specific *ACVRL1*^hi^ signature identified patients with an exceedingly poor survival in breast cancer. As a proof of concept, *in vivo* combination of ALK1 inhibition and immune checkpoint blockade resulted in significantly reduced metastatic disease. Finally, by means of 3D modeling of the immune-endothelial interface, FACS immunophenotyping, and multiplex imaging coupled with spatial analysis, we further uncovered the multimodal local and systemic effects of an impaired ALK1 signaling in the TME.

In conclusion, our work uncover that ALK1 embodies a dual angiogenic and immunomodulatory function in breast cancer, thereby providing a rationale for the re-evaluation of ALK1-blocking agents in combination with immune checkpoint blockade.

## Results

### Inhibition of ALK1 alters the extent of immune infiltrate in experimental primary and metastatic breast cancer

Based on our previous observations that *ACVRL1* expression is associated with immune features of the TME in a series of human solid malignancies (23), we confirmed the generality of this regulation in breast cancer. As expected, the network of genes correlated with *ACVRL1* expression paralleled a similar pattern in the TCGA Breast Invasive Carcinoma cohort (24, 25), with a highly significant enrichment for angiogenesis and hypoxia, in keeping with the reported endothelial expression of ALK1 (Table S1). However, the largest group of gene set enrichment analysis terms fell into a broad collection of processes defining the functionality and regulation of the immune cell compartment, *e.g.* interferon (IFN)-ɣ response, IL2-STAT5 signaling, and TNF-α signaling *via* NF-κB (Figure S1A and Table S1).

In light of these results, we immunostained experimental breast cancer tissue from transgenic MMTV-PyMT mice treated with ALK1-Fc (13) (Figure 1A), a decoy receptor that traps the high affinity ligands BMP-9 and BMP-10, resulting in a pruned vasculature and reduced tumor load. Assessment of DAB-IHC revealed an increased abundance of immunoreactivity against the broad leukocyte marker CD45 in ALK1-Fc-treated tumors compared with control IgG2a (Figure 1B). An analogous rise was observed for the T lymphocyte-restricted marker CD3 (Figure 1C). Furthermore, RNA was extracted from tissue sections from the same tumors and used as input for a qRT-PCR array focusing on immune-related genes. As shown in Figure 1D (and Table S2), tumors exposed to ALK1-Fc displayed a substantial increase in the expression of genes broadly regulating the identity and activation state of different immune cell types, including *Cd40lg, Cd4*, *Cd3d, Ifng*, *Il4*, and *Pdcd1lg2* (PD-L2).

**Figure 1.**
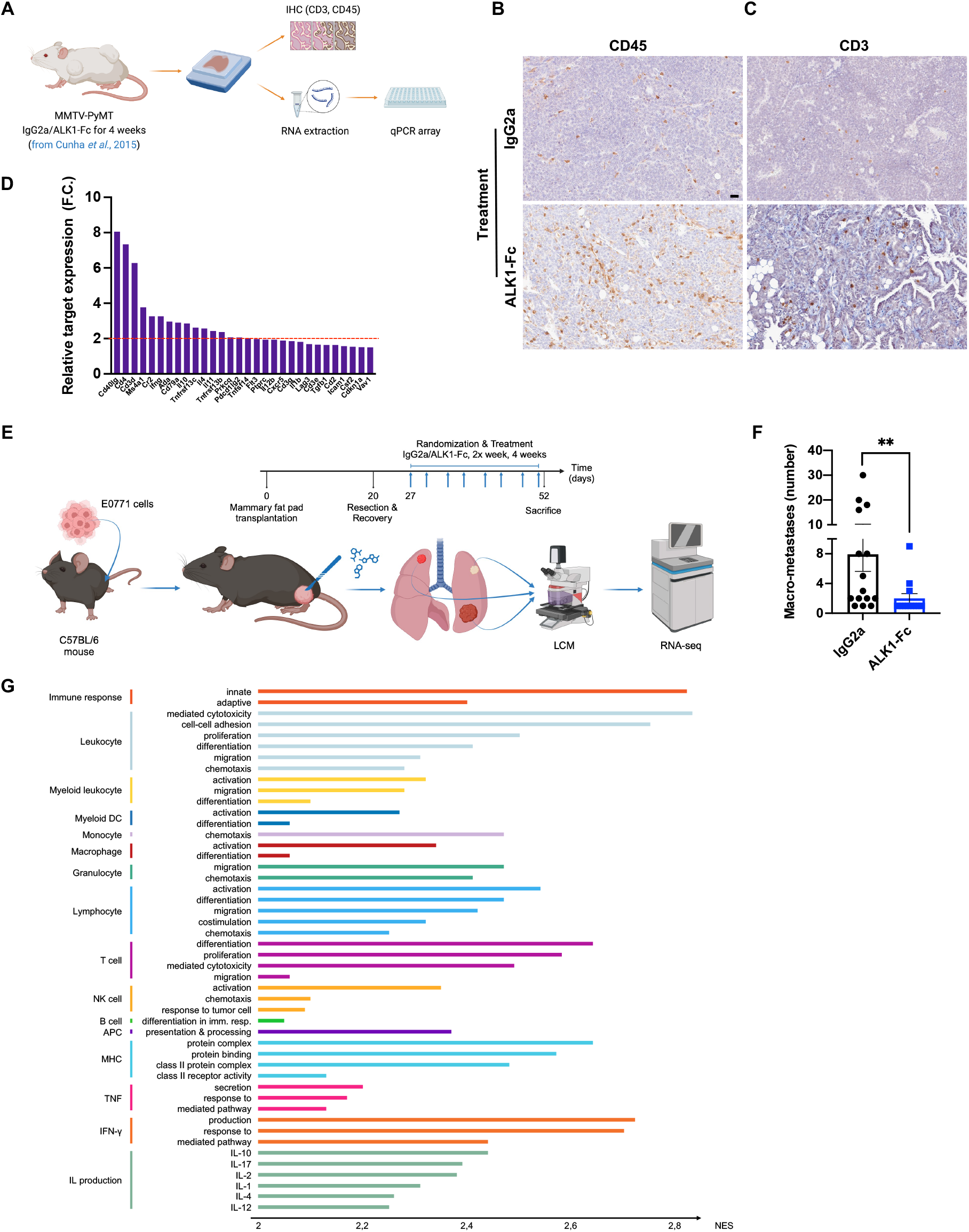
Inhibition of ALK1 alters the extend of immune infiltrate in experimental and metastatic breast cancer. (A): Analysis of archival breast cancer tissue from the transgenic MMTV-PyMT mouse model treated with IgG2a or ALK1-Fc (13). (B-C): Representative fields of DAB-IHC for CD45 (B), and CD3 (C), ALK1-Fc *vs* IgG2a. Scalebar: 100 μm. (D): Plot displaying the fold-change expression of target genes from the qRT-PCR, ALK1-Fc *vs* IgG2a. F.C.: Fold Change. (E): Experimental design of the adjuvant trial based on the orthotopic transplantation of 5×10^5^ E0771 cells in syngeneic C57BL/6 hosts. (F): Quantification of macro-metastases at sacrifice, ALK1-Fc *vs* IgG2a. Data displayed as mean with standard error of the mean (SEM). P-value: Mann-Whitney U-test. (G): Selection of significant gene ontology terms (adjusted p-value < 0.05) from the analysis performed on bulk RNA-sequencing of E0771 lung metastases. Normalized enrichment score (NES) values, ALK1-Fc *vs* IgG2a. DC: dendritic cell; APC: antigen presenting cell; MHC: major histocompatibility complex; IFN: interferon; TNF: tumor necrosis factor; IL: interleukin.

Primary tumors from MMTV-PyMT mice, or resulting from E0771 breast cancer cell transplantation, did not respond to either PD-1 or CTLA-4 inhibitors alone, or in combination with ALK1-Fc (Figures S1B-C). To better reflect the clinical management of breast cancer patients, and acknowledging the potent immunosuppressive effect of primary tumors (26), we next modeled an adjuvant therapy setup (Figure 1E). Following the engraftment of E0771 breast cancer cells in the abdominal mammary fat pad of syngeneic recipient hosts, incipient tumors were surgically resected when they reached 13 mm in the largest diameter. Five to 7 days after surgery, mice were randomized to receive IgG2a or ALK1-Fc for up to 4 weeks, while carefully monitoring for signs of disease spread to the lungs. At the experimental endpoint, administration of ALK1-Fc significantly reduced the number of macro-metastases per mouse, without affecting metastatic incidence (Figure 1F).

Based on the relative size and distribution of the metastases, lung lesions were either manually isolated or laser-capture microdissected, and further processed for bulk RNA sequencing. Enrichment analysis outlined an overrepresentation in tumors from ALK1-Fc-treated mice of pathways related to inflammatory response, complement, allograft rejection, as well as apoptosis, whereas DNA repair was greatly underscored (Figure S1D and Table S3). Strikingly, the statistically significant ontology terms covered both the innate and the adaptive arms of the defense response, encompassing more specialized functions such as interleukin production, response to interferon-ɣ, activation, migration, and chemotaxis of a series of distinct immune cell types (Figure 1G and Table S3).

Jointly, these data suggest that ALK1 inhibition promotes an inflammatory TME that may be further exploited for improved tumor control.

### Murine and human macrophages express *Acvrl1*/*ACVRL1*

The exclusive enrichment for immune processes following ALK1 inhibition made us wonder about the etiology of this regulation. Although an altered angiocrine signaling caused by inhibition of ALK1 is a logical and expected possibility considering the reported endothelial specificity of expression, it is tempting to speculate that a population of immune cells may directly react to ALK1 modulation, given the scale of the response observed at the gene expression level. To test the latter hypothesis, primary tumors from MMTV-PyMT mice were dissociated to single-cell suspensions, and expression of *Acvrl1* was queried by qRT-PCR of RNA isolated from immune cell populations sorted by flow cytometry (Figures 2A and S2A). Isolated endothelial cells and malignant cells served as positive and negative controls for *Acvrl1* expression, respectively (Figure 2B). Strikingly, as shown in Figure 2B, a subset of myeloid cells readily expressed *Acvrl1*, with Ly6C^-^ CD64^+^ macrophages, CD11b^+^ Ly6C^hi^ monocytes, CD11b^+^ MHCII^+^ dendritic cells, and CD11b^+^ Ly6G^-^ neutrophils displaying the highest expression levels. Additionally, the expression of *Acvrl1* bore functional implications, as stimulation of bone marrow (BM)-derived macrophages with the high affinity ligand BMP-9 significantly upregulated the expression of the downstream target gene *Id1* (Figure 2C).

**Figure 2.**
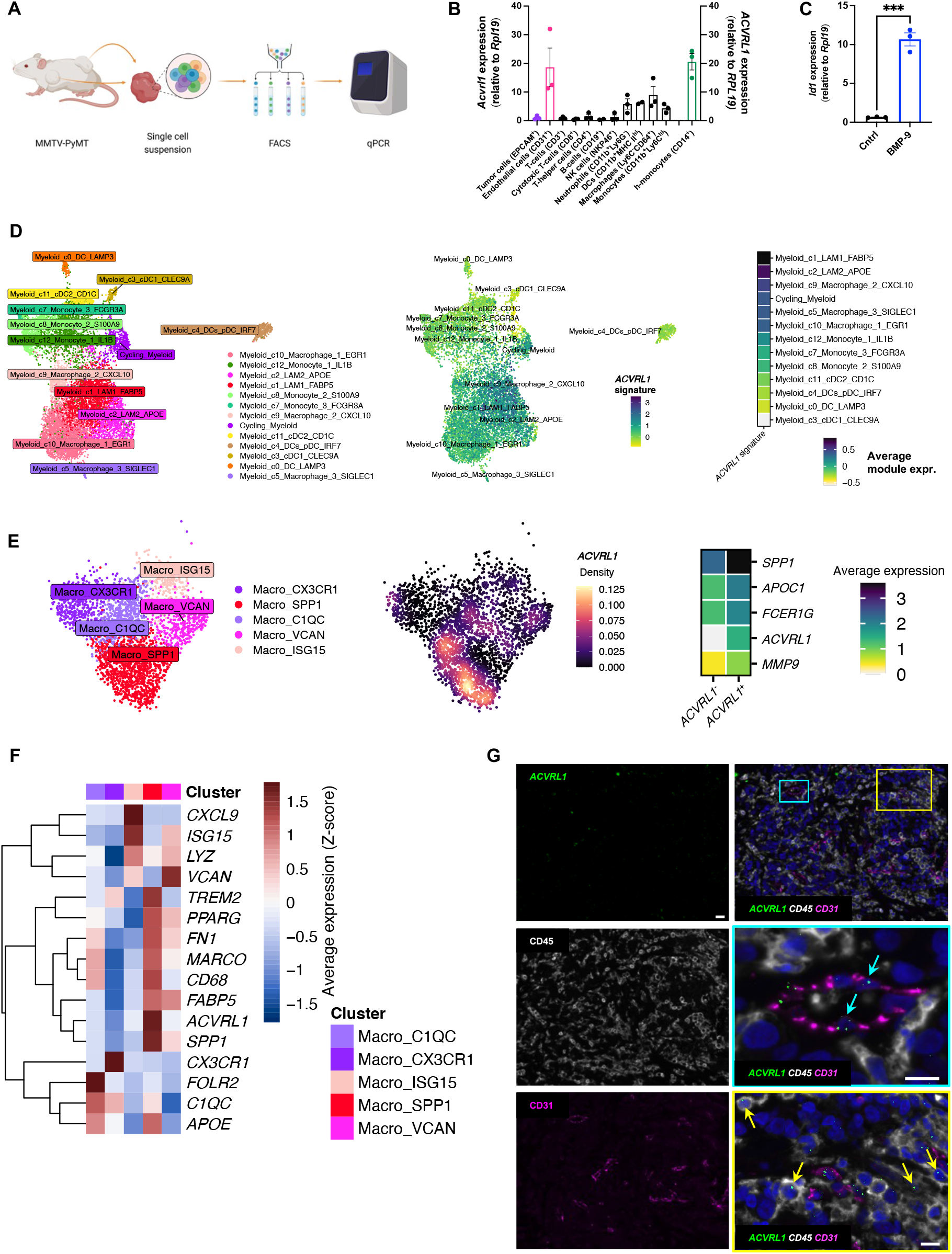
Murine and human macrophages express *Acvrl1*/*ACVRL1*. (A-B): Study design (A) for the quantification of *Acvrl1* expression in FACS-sorted immune cell populations from the MMTV-PyMT model (3 pooled experiments) (B). Positive (CD31^+^ endothelial cells) and negative (EpCAM^+^ epithelial cells) controls highlighted in magenta and purple, respectively. Expression of *ACVRL1* in freshly isolated human CD14^+^ monocytes from healthy donors (green). Data displayed as mean with SEM. (C): Expression of *Id1* in unstimulated (Control) *vs* BMP-9-stimulated bone marrow (BM)-derived macrophages (representative of 3 independent experiments). Data displayed as mean with SEM. p-value: unpaired, two-tailed t-test. (D): Overlay of a TAM-specific *ACVRL1* signature onto myeloid cells of a human breast cancer scRNA-seq dataset (32). (E): Expression of *ACVRL1* in TAMs from the scRNA-seq atlas of immune phenotypes (*29, 30*). Density plot of the expression of *ACVRL1* in TAMs. The average expression of the genes composing the TAM signature is presented in the heatmap for *ACVRL1*^+^ and *ACVRL1*^-^ TAM populations. (F): Heatmap of the expression of *ACVRL1*, cluster markers, and prototypical TAM-markers in the scRNA-seq atlas of immune phenotypes (29). (G): Dual RNAscope ISH coupled with mIHC in human breast cancer. The protein markers CD31 (magenta) and CD45 (white) were used to describe the cellular distribution of the *ACVRL1* probe (green). Scalebar 20 μm. Two inlets were annotated to highlight endothelial (cyan inlet/arrows) or immune-restricted accumulation of *ACVRL1* (yellow inlets/arrows). Inlet scalebar 10 μm.

Scrutiny of the recently published genomic catalogue of the adult human breast (27) consolidated the expression of *ACVRL1* in a series of annotated myeloid cell types (Figure S2B), including a variable yet conserved proportion (4-8%) of Macro -M2, -lipo and -IFN subsets as the fractions with the highest expression, indicating that *ACVRL1* is a feature of several macrophage phenotypes and states. Moreover, our assessment of the expression of *ACVRL1* by means of qRT-PCR revealed prominent mRNA expression in freshly isolated human CD14^+^ monocytes from the peripheral blood of healthy donors (Figure 2B). In agreement with this finding, *ACVRL1* was also discernible in CD14^+^ monocytes in a single-cell (sc)RNA-seq collection of the human bone marrow (*28*) (Figure S2C).

In the context of cancer, we started off by generating and validating a TAM-specific *ACVRL1* signature. Significantly differentially expressed genes (DEGs) between *ACVRL1*^+^ and *ACVRL1*^-^ macrophages were extracted from a breast-specific map of immune phenotypes (29) with an updated cellular annotation (30) (Table S4). This list was further filtered through a triple-negative breast cancer (TNBC) scRNA-seq dataset (31) by applying a stringent criteria selection (*i.e.* high expression restricted to the macrophage cluster; Table S5), leading to four genes: *SPP1*, *APOC1*, *FCER1G*, and *MMP9*. The final signature –which included these four genes as well as *ACVRL1*— teased out a distinct cluster of myeloid cells and TAMs when imposed on two additional breast cancer scRNA-seq metadata (31, 32) (Figures S2D-E), with the highest average signature expression in lipid-associated macrophages (LAMs; Figure 2D). Next, we queried the approximately 400 DEGs between *ACVRL1*^+^ *vs ACVRL1*^-^ TAMs in relation to the hallmarks of intratumoral heterogeneity that were recently mapped out (33). Within the macrophage cell type, this analysis confirmed a significant enrichment for lipid-related, glycolysis, and proteasomal degradation meta-programs (Figure S2F and Table S6).

When closing in on TAMs, inspection of the breast-restricted immune atlas revealed the highest expression of *ACVRL1* in angiogenesis-associated, pro-tumorigenic *SPP1*^+^ TAMs (34) (Figures 2E and S2G). Reassuringly, the average expression of the genes included in the signature was higher in *ACVRL1*^+^ vs *ACVRL1*^-^ TAMs (Figure 2E). Within the *SPP1*^+^ cluster, *ACVRL1* expression strongly mirrored other immunosuppressive markers, such as *FABP5* and *TREM2,* with both genes further converging into the previously reported LAM phenotype (32, 35) (Figure 2F). Notably, *TREM2* also denotes immunosuppressive (and M2-like) TAMs with a putative monocytic origin (36, 37). To expand on the predicted recruited origin (and supported by *ACVRL1* expression in CD14^+^ monocytes), our analysis highlighted the distinct expression pattern of *ACVRL1* relative to *FOLR2,* which unequivocally identifies resident APOE^+^ TAMs (38); indeed, *FOLR2* expression exclusively segregated in the *C1QC*^+^ TAM cluster (Figure 2F).

Finally, to validate these results in patient material, the expression of ALK1 was probed in human breast cancer specimens by RNAscope (due to the paucity of antibodies specific for ALK1 suitable for immunostaining) coupled with highly sensitive multiplexed immunohistochemistry (mIHC) to identify constituent cell types. As expected, the signal of *ACVRL1* readily overlapped with the endothelial marker CD31 (cyan inlet/arrows, Figure 2G). Moreover, the characteristic dots of the RNA-ISH clearly accumulated in CD45-positive cells in the tissue (yellow inlet/arrows).

Taken together, our data reveal that expression of ALK1 is not, as previously reported, exclusive for the endothelium, and that a population of recruited TAMs is also characterized by *ACVRL1* expression across human breast malignancies.

### *ACVRL1*-expressing TAMs display an immunosuppressive phenotype associated with resistance to therapy and poor survival

To gain additional insight into the translational relevance of the molecular cues instigated by ALK1 in macrophages, we estimated survival outcomes in 1097 breast cancer patients from the TCGA repository (39) through Kaplan-Meier survival fractions, as well as a Cox proportional hazard model. After adjusting for stage, age at diagnosis, and estrogen receptor (ER) status, the *ACVRL1* signature*^high^* group exhibited worse disease-specific survival (DSS; p-value = 0.017, HR = 1.86 (95% CI = 1.14-3.03); Figure 3A), as well as progression-free interval, (PFI; p-value = 0.014, HR = 1.66 (95% CI = 1-16-2.38); Figure S3A), compared to the group of patients with the lowest *ACVRL1* signature expression; a similar trend was observed for overall survival (OS; p-value = 0.19, HR = 1.23 (95% CI 0.87-1.75); Figure S3B). These results were validated in the METABRIC (40) dataset, which confirmed the significance for DSS (p-value = 0.018, HR = 1.34 (95% CI= 1.11-1.60; Figure 3B), and the trend for OS (p-value = 0.077, HR = 1.14 (95% CI = 0.99-1.31); Figure S3C). Moreover, the negative impact on survival of a high expression of the *ACVRL1* signature is in line with the previously reported prognostic value of pro-tumorigenic *SPP1*^+^ TAMs (29, 34).

**Figure 3.**
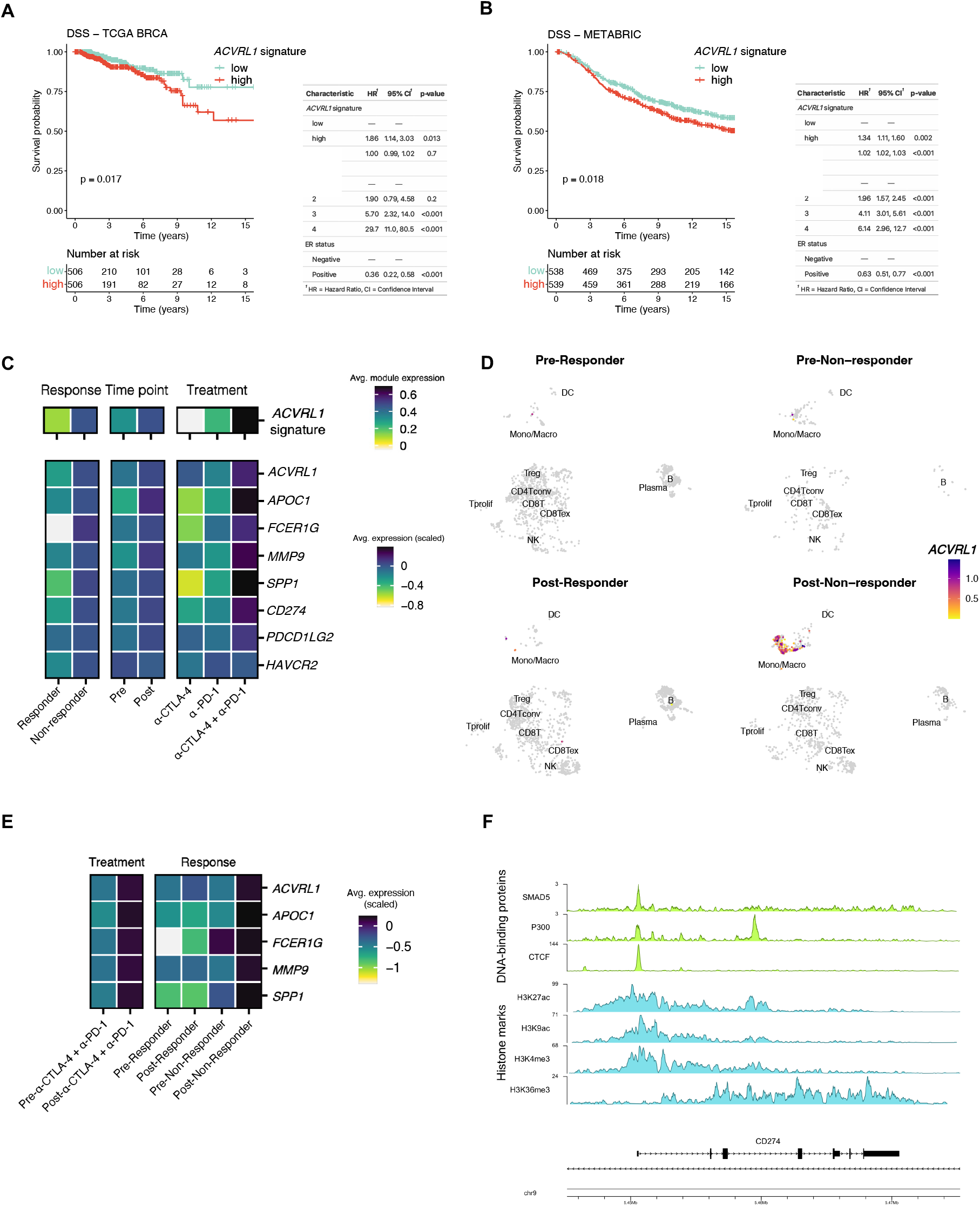
ACVRL1-expressing TAMs display an immunosuppressive phenotype associated with resistance to therapy and poor survival. (A-B): Survival analysis in the TCGA BRCA (39) (A) and METABRIC (40) (B) datasets. Patients were stratified into two risk groups based on the median value of the mean expression of a TAM-specific *ACVRL1* signature. The Kaplan-Meier curves show the disease-specific survival (DSS) probabilities of the high (red) and low (green) signature expression groups in the two cohorts. P-value: log-rank test. The tables summarize the relative Cox proportional hazard model analysis for each cohort. (C-E): Expression of *ACVRL1* in a CD45^+^-restricted scRNA-seq compendium of 48 melanoma patients treated with immune checkpoint inhibitors (42). The average expression of the 5-gene signature, and the average scaled expression of the individual genes are presented in a heatmap (C) based on response, time point, and treatment arm. The average expression of *ACVRL1* in the combined CTLA-4 and PD-1 inhibition group was imposed on the UMAP, and further split to create four different groups: pre-responder, post-responder, pre-non-responder, and post-non-responder (D). Contingent on the aggregated data points in (D), the average scaled expression of the five genes comprised in the *ACVRL1* signature is presented in a heatmap (E). (F): ChIP-seq plot from human hematopoietic tissue (43, 44). Visualization of DNA-binding proteins and histone marks in the promoter region of *CD274*.

Prompted by these results, we set out to determine how myeloid expression of ALK1 tied in with the clinical performance of immunotherapy. Limited by the inherent lack of extensive implementation in breast cancer (41), we resorted to melanoma, where the use of immune checkpoint inhibitors has revolutionized the clinical management of patients. Thus, we screened a cohort of 48 melanoma patients that were longitudinally sampled throughout therapy (42), in this case a regimen of anti-CTLA-4, anti-PD-1, or combined CTLA-4 and PD-1 blockade. The analysis of this dataset confirmed that ALK1-expressing macrophages represented a minor subset of TAMs at baseline (*circa* 7%), which were efficiently removed by therapy in the responder group (Figure S3D). Strikingly, ALK1^+^ TAMs emerged prominently upon treatment resistance, where they represented approximately 18 % of the tumor monocyte/macrophage cluster in the non-responder group (Figure S3E). Accordingly, the expression of the *ACVRL1* signature –as well as the individual genes included in it—were significantly higher in non-responders *vs* responders, and in post-*vs* pre-treatment data points (Figures 3C and S3E). Notably, the highest frequency and average expression of *ACVRL1* were detected in the treatment arm combining dual CTLA-4 and PD-1 blockade. In light of this specific modulation, we homed in on the combined anti-CTLA-4 and anti-PD-1 group, and the data from this treatment arm was dichotomized based on response at each timepoint. As shown in Figures 3D-E, the *ACVRL1*^+^ myeloid cells present at baseline persisted throughout therapy, and further expanded in the non-responders. This regulation was even more striking when considering the *ACVRL1* signature, which showed an inherent difference already at baseline (Figure S3E), suggesting that components of this signature might be responsible for intrinsic, primary resistance to immunotherapy. Moreover, *ACVRL1* expression was tied to that of immunosuppressive markers like *CD274* (PD-L1) and *PDCD1LG2* (PD-L2) in non-responders in the global cohort (Figure 3C). These data imply that ALK1 signaling may be directly involved in the specification of an immunosuppressive phenotype in TAMs. In agreement with this hypothesis, a distinct peak for SMAD5, the signaling mediator downstream of ALK1, could be extrapolated in the promoter region of *CD274*/PD-L1 from ChIP-seq data of hematopoietic tissue in the ENCODE database (43, 44), further intersecting with a domain of euchromatin, indicative of direct accessibility to transcription factors (Figure 3F). This analysis revealed the specificity of the signal for SMAD5, as the DNA-binding and the histone mark patterns could not be replicated for *PDCD1LG2*/PD-L2 nor *HAVCR2*/TIM-3 (Figures S3F-G).

Collectively, these results offered us a rationale to combine ALK1 blockade with immune checkpoint inhibitors.

### Inhibition of ALK1 potentiates immunotherapy

Motivated by the previous results indicating an ALK1-dependent specification of a pro-tumorigenic immune state, we yet again exploited our resection-based adjuvant therapy pipeline, this time using the syngeneic 4T1 mammary carcinoma cell line. Unlike the E0771-based model, in which only 40% of the mice generated macro-metastases, the 4T1 cell line gives rise to tumors resembling stage IV human TNBC with a complete penetrance to different metastatic sites in a time-dependent manner. In an initial trial modeling a pre-metastatic setting, treatment with ALK1-Fc only showed a trend towards controlling disease progression, possibly reflecting the more aggressive nature of the 4T1 cell line (Figures S4A-B). Next, in a more advanced stage experimental setup, mice were randomized to receive post-surgery therapy with control IgG2a, ALK1-Fc, immunotherapy (IT) comprising dual PD-1 and CTLA-4 inhibition, or a combination of ALK1-Fc and IT (Figure 4A). At sacrifice, the total lung weight was recorded as a readout for metastatic burden. Notably, the cohort of mice that received combined treatment with ALK1-Fc and IT exhibited the lowest metastatic load of all groups, with a significantly reduced lung weight compared to immune checkpoint blockade alone (Figure 4B).

**Figure 4.**
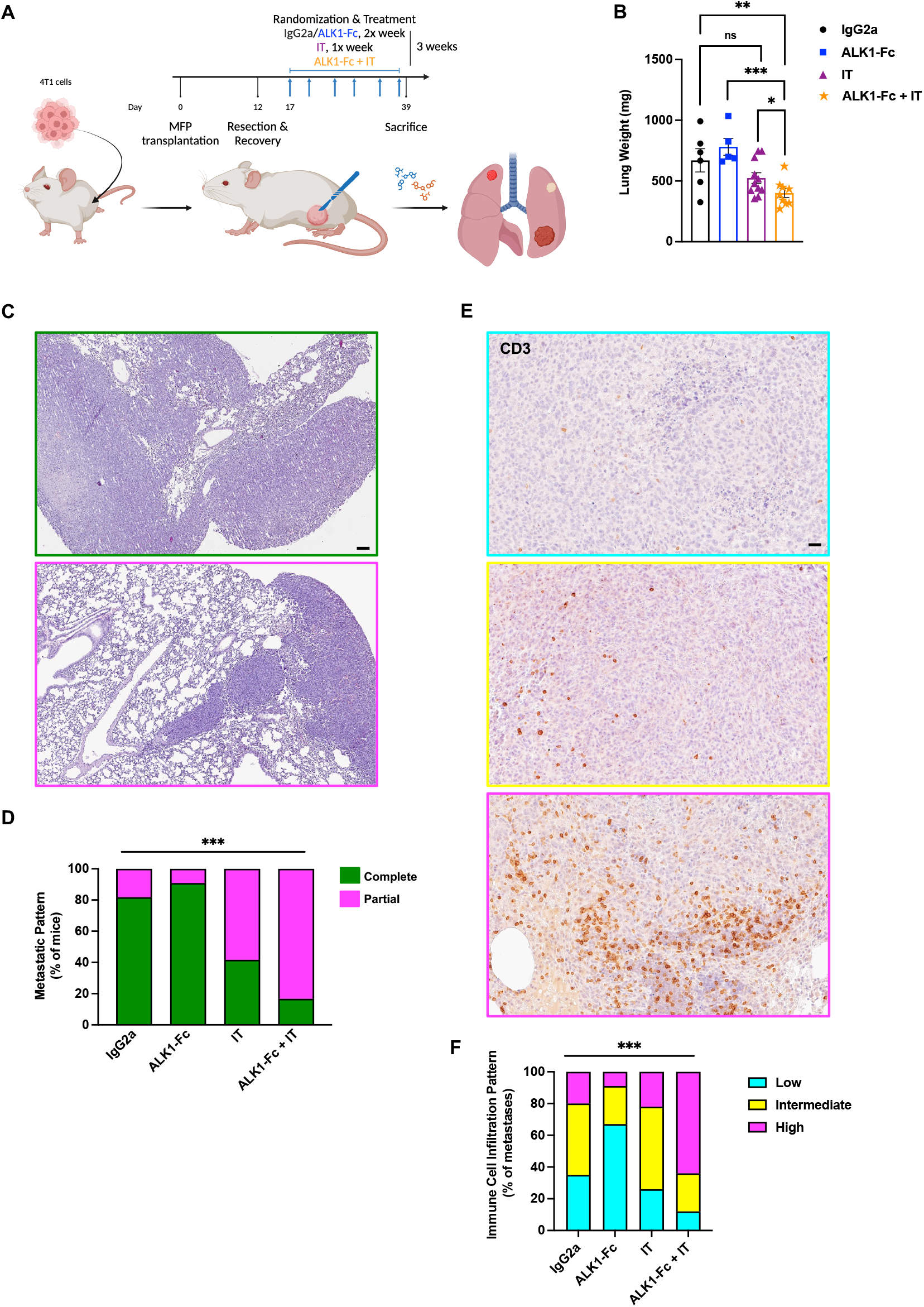
Inhibition of ALK1 potentiates immunotherapy. (A-B): Experimental design of the adjuvant trial based on the orthotopic transplantation of 5×10^4^ 4T1 cells in syngeneic BALB/c hosts (A). Immunotherapy (IT) consists of a dual inhibition of PD-1 and CTLA-4. MFP: mammary fat pad. Quantification of total lung weight as a proxy for metastatic load (B). Data displayed as mean with SEM. p-value: unpaired, two-tailed t-test (ns: not significant). (C-D): Hematoxylin and Eosin staining of whole lung sections from the different cohorts in the 4T1 adjuvant trial. Representative pictograms of whole lung metastatic infiltration (green; C) or partial metastatic outgrowth (magenta), quantified in (D). Scalebar: 100 μm. p-value: χ2 test. (E-F): DAB-IHC for CD3 in whole lung sections from the different cohorts (E), and quantification of the proportions of the CD3 distribution in the metastases (F). The staining pattern was categorized in low (cyan), medium (yellow), and high (magenta). Scalebar: 50 μm. p-value: χ2 test.

We corroborated these results by assessing the lung parenchyma in the different cohorts. More than 80% of the mice treated with either IgG2a or ALK1-Fc bore metastases that virtually invaded the whole lungs (green; Figures 4C-D). Conversely, approximately 60% of the IT group and 85% of the animals in the ALK1-Fc + IT group exhibited distinct malignant lesions that were clearly encapsulated within adjacent/normal lung tissue (magenta; Figures 4C-D). In line with this, evaluation of the distribution of CD3 staining within metastatic lesions supported that control tumors were more generally immune-excluded, and that they became increasingly more inflamed and readily infiltrated by lymphocytes as IT or a combination of ALK1-Fc + IT were administered (Figures 4E-F). Even though processing of bulk RNA sequencing did not reveal major transcriptional differences between the cohorts, a series of specific genes with a clear involvement in angiogenesis and immune cell modulation, including *Dysf*, *Ly6g2*, *Ccl8*, and *Lrrc32*, were significantly up- or down-regulated in the combination treatment group *vs* either monotherapy (Figures S4C-E and Table S7).

### Inhibition of ALK1 elicits local and systemic effects on the immune landscape

Next, we sought to define which immune cell type is responsible for the integration of the signals coming from the different inhibitory cues during anti-angiogenic immunotherapy. For this purpose, resected 4T1 primary tumors, lung metastases and peripheral blood from a second equivalent experiment were used as input for a multicolor fluorescence-activated cell sorting (FACS) immunophenotyping approach devised to dissect the lymphoid and myeloid arms of the immune system (45) (Figures 5A-B). Combined treatment with ALK1-Fc and immune checkpoint blockade directly acted on monocytes and macrophages in the metastatic tissue, with a significant increased frequency of Ly6C^hi^ CD64^-^, and a concomitant reduction of Ly6C^hi^ CD64^+^ cells (Figures 5C-D). In line with this, the combined treatment also significantly lowered the proportion of CD64^+^ macrophages in the lungs (Figure 5E), and increased several subsets of dendritic cells (Figures 5F-H). In light of the observed expression pattern of ALK1, the combined treatment indirectly altered the lymphoid landscape (Figures S5A-D) by significantly increasing the frequency of NK cells (Fig. 5I), CD3^+^ T-cells (Fig. 5J), CD4^+^ T-helper (Fig. 5K), and a trend for CD8^+^ cytotoxic T-lymphocytes (CTLs; Figure 5L). Moreover, these intra-metastatic changes were coupled with systemic alterations in peripheral blood composition (Figures S5E-I), with ALK1-Fc + IT remarkably stalling the differentiation from CD64^-^ to CD64^+^ monocytes (Figures 5M-O). This effect appeared to be reliant on ALK1 signaling, as the same modality of regulation was also detected in the ALK1-Fc cohort.

**Figure 5:**
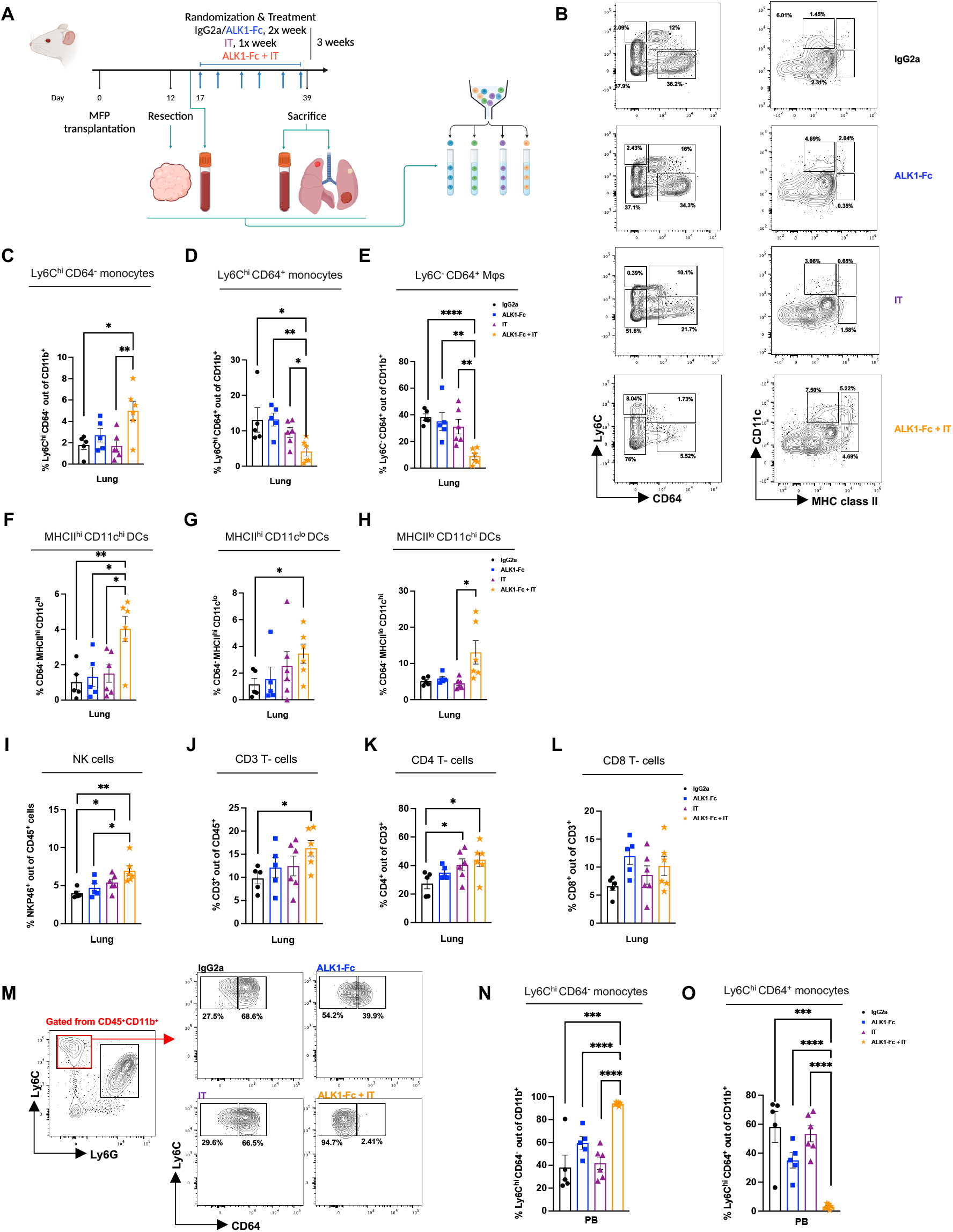
Antiangiogenic immunotherapy elicits direct and indirect effects on the immune landscape. (A): Experimental design of the 4T1-based adjuvant trial to study different population of myeloid and lymphoid cells via FACS. (B): representative purity check plots for the myeloid compartment in the different cohorts. (C-D): From the CD11b^+^ gating, relative abundance of monocytes in lung tissue: Ly6C^hi^ CD64^-^ (C), and Ly6C^hi^ CD64^+^ (D). Data displayed as mean with SEM. p-value: unpaired, two-tailed t-test. (E): From the CD11b^+^ gating, relative abundance of Ly6C^-^ CD64^+^ macrophages in lung tissue. (F-H): From the CD64^-^ gating, relative frequency of dendritic cells in lung tissue: MHCII^hi^ CD11C^hi^ (F), MHCII^hi^ CD11C^lo^ (G), and MHCII^lo^ CD11C^hi^ (G). Data displayed as mean with SEM. p-value: unpaired, two-tailed t-test. (I-L): From the CD45^+^ population, relative frequency of NKP46^+^ NK cells (I), CD3^+^ T-cells (J), CD4^+^ T-helper cells (K), and CD8^+^ cytotoxic T lymphocytes (L). Data displayed as mean with SEM. p-value: unpaired, two-tailed t-test. (M-O): From the CD45^+^ CD11b^+^ cells, purity check of circulating monocytes in peripheral blood (M). Relative frequency of circulating monocytes: Ly6C^hi^ CD64^-^ (N), and Ly6C^hi^ CD64^+^ (O). Data displayed as mean with SEM. p-value: unpaired, two-tailed t-test.

Altogether, these data suggest a dynamic regulation at multiple levels in the hematopoietic differentiation cascade and the metastatic immune landscape by the addition of ALK1-Fc to an immunotherapy regimen.

### Vascular immune features reflect differential response to anti-angiogenic immunotherapy

To combine the data coming from the necropsy observations and the immunophenotyping, we developed a customized mIHC panel to delineate the spatial relationships of endothelial cells and a series of immune cell types (Figure 6A and Table S8). In the absence of reliable reagents for the specific detection of ALK1, and to clearly distinguish the presence of different cell types, we selected CD31 to mark endothelial cells, and TREM2 as a proxy to identify pro-tumorigenic ALK1^+^ TAMs. In addition, we stained for tumor infiltrating lymphocytes (TILs): CD4^+^ T-helper cells, CD8a^+^ CTLs, and B220^+^ B-cells. Finally, (tumor) epithelial cells were distinguished *via* EpCAM. Following segmentation to classify metastatic tissue and normal/adjacent lung (Figure 6B), cell identification and phenotyping (Figures 6C-D) mirrored the features captured with FACS, IHC, and RNA-seq. Given the high level of heterogeneity within and between lesions, we focused on the cohort exposed to combined ALK1-Fc and immune checkpoint blockade, in which we observed a phenotypic range reflecting the overall response surveyed at sacrifice (Figure 6E), as well as for human patients undergoing immunotherapy. Metastatic tissue from a mouse that responded to therapy (based on the lung weight and metastatic count at sacrifice, as well as the H&E histology) encompassed the lowest abundance of CD31^+^ endothelial cells and TREM2^+^ TAMs (Figure 6E; responder). In contrast, a stable vasculature paralleled by an increased TAM-infiltration characterized the lesion of a mouse that had a controlled metastatic burden but relapsed at the primary site (Figure 6E; partial). Finally, prominent metastatic lesion vascularization and a much higher proportion of CD4^+^ cells within TILs epitomized the group of mice that did not respond/progressed upon combined therapy administration (Figure 6E; non-responder). This distribution was even more evident when considering the spatial density (Figure 6F): in this analysis, the responder lesion in the combination treatment group displayed the highest density of both T effector and T helper lymphocytes, together with the lowest density of endothelial cells and TAMs. The segmentation further captured that the influx of effector T cells was specific for the metastatic tissue, as the density of the CD8^+^ CTLs was comparable in the lung segments across the samples. In keeping with this, the CTL-to-macrophage ratio promptly correlated with response (Figure 6G).

**Figure 6:**
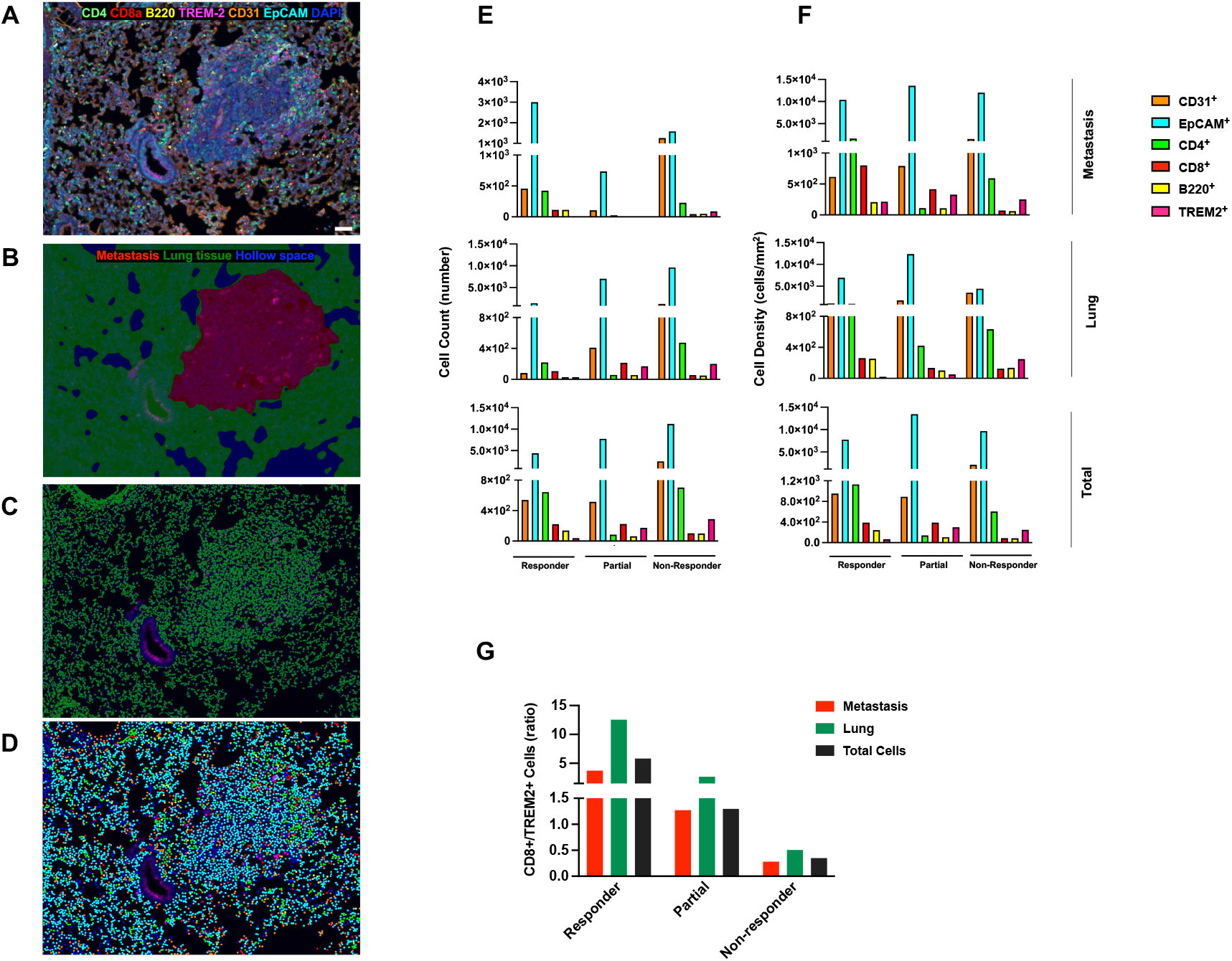
Vascular immune features reflect differential response to antiangiogenic immunotherapy. (A): Customized multiplexed IHC (mIHC) staining of 4T1 metastases to the lungs. An antibody panel was developed to detect CD31^+^ endothelial cells (orange), EpCAM^+^ epithelial cells (both lung epithelium and breast cancer cells, cyan), TILs (CD4^+^ T helper, green; CD8a^+^ cytotoxic T lymphocytes, red; B220^+^ B-cells, yellow), and TREM2^+^ recruited TAMs (magenta). Scalebar 50 μm. (B-D): A machine learning-based algorithm was trained to discriminate metastatic tissue (red) from lung and hollow space (green and blue, respectively; B), followed by cell segmentation (C). Phenotyping (D) is visualized as a dot with the same color coding as in (B). Cells (based on DAPI detection) negative for any of the markers included in the antibody panel are displayed in blue. (E-G): total cell counts per phenotype (E), and cell density per phenotype (cells/mm^2^; F) within the different tissue segments. The CD8^+^ T-cells/TREM2^+^ TAMs ratio (G) was calculated from cell densities.

Taken together, spatial analysis of the cellular composition of responding or resistant lung metastatic lesions revealed that the combination of ALK1-Fc with immunotherapy promoted an active anti-tumor environment, while preserving sensitivity to the anti-angiogenic therapy.

### ALK1 affects the hematopoietic progenitor cell niche in the bone marrow

Remodeling of the metastatic TME based on the response to ALK1-Fc and immunotherapy also corroborated the known and expected ALK1-dependent effects on the vasculature. Thus, we next sought to clarify whether endothelial ALK1 modulation limited the trans-endothelial migration of myeloid cells, as an explanation for the observed paucity of immunosuppressive TAMs in the metastatic milieu following ALK1 inhibition. To this end, murine lung endothelial cells (mLECs) engineered to express two different shRNA constructs against *Acvrl1* (shA07 and shA09; Figure S6A) or a scrambled control construct (shCtrl) were seeded to populate the top channel of a 3-lane microfluidic organ-on-a-chip device (Fig. 7A). A barrier integrity test confirmed the generation of an intact 3D vascular tube, allowing its perfusion with macrophages (Figure S6B and Video 1). In turn, the CCL2 chemoattractant in the bottom lane stimulated the macrophages to migrate to the intermediate channel. Laser scanning confocal microscopy revealed a consistent trend towards a curtailed migration through *Acvrl1*-silenced endothelium across our longitudinal imaging schedule (Figure 7B), with a significant difference for the shA09 at 24 hours, indicating that ALK1 indeed regulates extravasation of monocytes.

**Figure 7.**
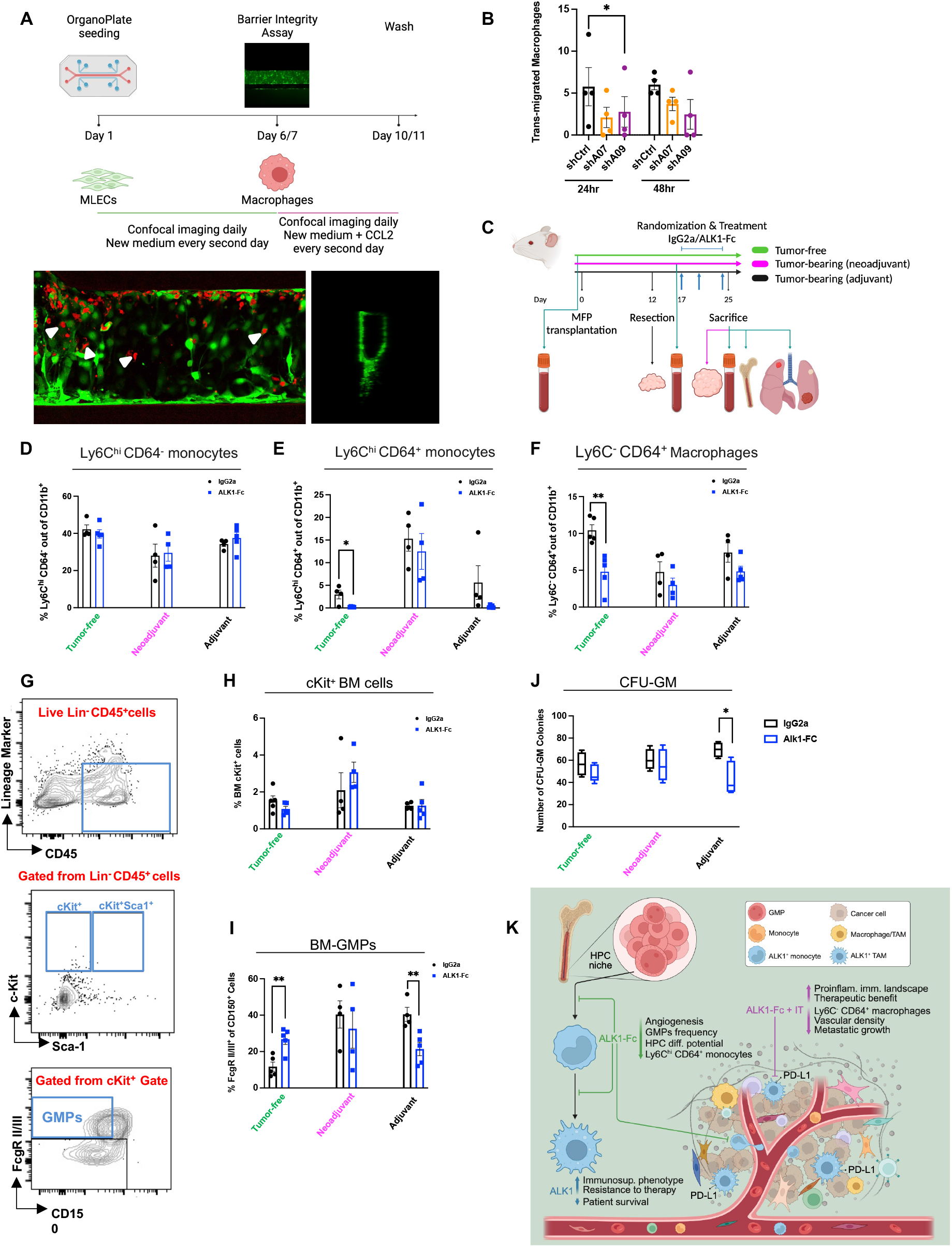
ALK1 affects the hematopoietic progenitor cell niche in the bone marrow. (A): Experimental design based on the Organ-on-a-chip assay with 3 channels. Representative images of a longitudinal and a cross section of the 3D tube are presented in the bottom panel. (B): Quantification of macrophage transendothelial migration at 24 and 48 hours. Data displayed as mean with SEM. p-value: unpaired, two-tailed paired t-test. (C-E): Experimental design of the short-term trial based on the orthotopic transplantation of 5×10^4^ 4T1 cells in syngeneic BALB/c hosts (C). Frequency of circulating Ly6C^hi^ CD64^-^ (D) and CD64^+^ (E) monocytes in peripheral blood. Data displayed as mean with SEM. p-value: unpaired, two-tailed t-test. (F): Frequency of Ly6C^-^ CD64^+^ macrophages in lungs. Data displayed as mean with SEM. P-value: unpaired, two-tailed t-test. (G-I): FACS plot and gating strategy of c-Kit^+^Lin^-^Sca1^-^ progenitor cells (46) extracted from the bone marrow (G). GMP: granulocyte-macrophage progenitor. Frequency of c-Kit^+^ (H) and GMP (I) cells from the bone marrow extracts. Data displayed as mean with SEM. P-value: unpaired, two-tailed t-test. (J): Quantification of colony formation plating efficiency of c-Kit-enriched bone marrow cells. Data displayed as mean with SEM. P-value: paired, two-tailed t-test. (K): summary.

Subsequently, given the modulation of circulating monocyte populations observed following adjuvant ALK1 inhibition (Figures 5N-O), we investigated whether ALK1 signaling acted upstream of macrophage functional states. We set up an *in vivo* experiment based on the 4T1 resection model to monitor the circulating monocytes in the framework of a short-term pharmacological inhibition of ALK1 (Figure 7C). Based on Ly6C and CD64 gating in flow cytometry analysis, we determined that ALK1-Fc reduced the frequency of circulating Ly6C^hi^ CD64^+^ monocytes in tumor-free mice, and produced a similar trend in the adjuvant setting (Figure 7D-E and S6C). Of note, no effect was appreciable in the neoadjuvant setting, reinforcing the vital role of primary tumor-derived immunosuppressive signals and crucial differences between the primary tumor ecosystem and metastatic lesions (26). Similarly, infiltration of CD64^+^ macrophages was significantly blunted in tumor-free lungs of ALK1-Fc-treated mice compared to the IgG2a group (Figure 7F), indicating that mobilization of monocytes is under the control of ALK1 activity and independent of altered angiogenic stimuli emanating from the tumor.

Taking these results into account, we next sought to clarify which step of the hematopoietic cascade is reliant on ALK1. Bone marrow cell evaluation (46) from the experimental groups showed a significant reduction in the population corresponding to the granulocyte-macrophage progenitors (GMP) in the adjuvant ALK1-Fc group, while there was no significant change in the proportion of c-Kit^+^ progenitor cells (Figure 7G-I). These results are evocative of the pattern evinced from the bone marrow atlas (28) (Figure S2C), which singled out the highest average expression of *ACVRL1* in the GMP cluster. Finally, in agreement with the reduction of BM-GMPs, we also observed a significant decrease in the potential of BM-derived progenitors to differentiate *ex vivo* into colony-forming unit-granulocyte-macrophage (CFU-GM)-colonies following exposure to adjuvant ALK1-Fc treatment *in vivo* (Figure 7J).

In conclusion, our data demonstrate that ALK1 functionally regulates the microenvironment in two specific compartments: locally, by pruning angiogenesis and fine-tuning permissiveness to monocyte/macrophage trans-endothelial migration; and systemically, by determining the breadth of monocyte differentiation and mobilization in the bone marrow.

## Discussion

The data presented in our work delineate a novel role for the hitherto endothelium-restricted ALK1 receptor. Through a series of *in vivo* models, clinically relevant treatment schedules, mechanistic studies, and integration of *in silico* datasets, we uncover and validate the retained expression of ALK1 in a population of CD14^+^ monocytes in physiological and pathological breast. When zooming in on cancer, these monocytes give rise to immunosuppressive TAMs that support disease progression upon failure of therapeutic regimens. We provide compelling evidence that the monocyte-to-TAM differentiation is the final step in a cascade of events that spring from the HPCs niche in the bone marrow, where ALK1 defines the potential of early progenitor cells (Figure 7K).

Pharmacological blockade of endothelial ALK1 signaling gained considerable attention in the mid-2010s following a series of encouraging pre-clinical and clinical trials in a range of different tumor types. Given the more restricted pattern of expression compared with the ubiquitous VEGF receptor 2 (VEGFR2), ALK1 inhibition became an enticing alternative to the promise of the “anti-angiogenic revolution” that ensued with the initial approval of bevacizumab. Despite this, the development of the ALK1 ligand trap dalantercept was blocked due to the lack of additional benefits, raising the issue of the initial patient selection as well as the optimal therapeutic combination partner. In the absence of biomarkers for ALK1 status, the tumor types selected for escalation studies and phase 2 trials failed to offer much ground for the anti-tumor activity of dalantercept, even more so when combined with standard-of-care agents that were already suboptimal in terms of efficacy. As we were unable to access samples collected during such clinical trials, the initial indication of an active involvement of tumor immunity stems from the retrospective analysis of archival breast cancer tissue from MMTV-PyMT mice, in which neoadjuvant ALK1-Fc monotherapy promotes a pro-inflammatory environment, with a markedly increased expression of *Cd3*, *Cd4*, as well as factors like *Ifng* (encoding for interferon-γ) and *Cd40l*. Collectively, these markers indicate a broad rewiring of the immune cell composition and activation status as a consequence of therapeutic pressure. For example, the CD40/CD40L axis is related to B-cell priming by CD4^+^ T-cells, and several agonists for CD40 have surfaced for clinical validation, *e.g.* mitazalimab and selicrelumab.

However, addition of immunotherapy to ALK1 blockade with a neoadjuvant regimen did not translate to better tumor control in the transgenic setting. This outcome corroborates akin published observations of combined VEGFR2 inhibition and immunotherapy (47), and it is in agreement with the generally accepted notion that immune checkpoint blockade is most efficient in tumors with high(er) mutational burden. Nonetheless, it is worth to mention that we observed a trend towards fewer malignant foci to the lungs in any treatment group that included neoadjuvant ALK1-Fc in our mouse models (Figure S1), suggesting that the molecular cues in the metastatic colonization and growth are somewhat different from those of the primary tumor mass.

The ability of ALK1 to directly specify an immunosuppressive phenotype of TAMs by regulating the transcription of *CD274* endows an unprecedented opportunity to investigate additional partners that could help sustain these immunomodulatory properties. Our proof-of-concept trial unequivocally indicates PD-1 and CTLA-4 inhibition as actionable therapeutic companions to ALK1 inhibition. An additional line of evidence that supports this combination comes from a clinical trial promoted by the pharmaceutical company Kintor, which resumed the development of GT-90001, a monoclonal antibody against ALK1 originally pursued by Pfizer. This early phase investigation determined the recommended dose of GT-90001 together with the anti-PD-1 nivolumab in advanced hepatocellular carcinoma (48). Except for thrombocytopenia, this combination showed a good safety profile, tolerability, and anti-tumor activity to justify further investigation.

The yield brought in by combined blockade of ALK1 and immune checkpoints reflects an intra-metastatic, as well as a systemic, underrepresentation of an immunosuppressive arm of the immune compartment, culminating with a depletion of recruited TAMs and circulating monocyte-derived cells, respectively. Even our bulk RNA-seq captures this macrophage-centered effect, as ALK1-Fc downregulates interferon-induced transmembrane (IFITM) 6 transcripts. The IFITM family of proteins is involved in cell-cell adhesion and cell differentiation; IFITM6 is specifically expressed in bone marrow-derived macrophages and its expression is increased in tumor-bearing mice (49). Moreover, despite the more aggressive growth pattern and metastatic penetrance of the 4T1-based platform, the addition of immunotherapy to ALK1 inhibition specifies for a significantly higher frequency of NK cells, CD4^+^ T-cells, as well as dendritic cells. Globally, our *in vivo* approaches confirm a boosted infiltration and functionality of T helper, CTLs, B-cells, and professional antigen-presenting cells following exposure to ALK1-Fc-encompassing therapies. These cell types are the major constituents of tertiary lymphoid structures (TLS), which have been characterized as prime determinants of sustained anti-tumor activity in patients treated with immunotherapy (50). Although protocols for the detection and spatial characterization of TLS have been standardized and widely available for clinical investigation, translation to murine models is not trivial, as these agglomerates cannot be generally observed in full (51). It becomes even more cumbersome in the metastatic setting, which suffers from the limited availability of data from the clinical experience (52). In agreement with this, the full-slide mIHC panel we customized for breast dissemination to the lungs did not reveal the presence of TLS in the metastases or embedded within the adjacent lung tissue, except for one single lesion displaying an interesting germinal center-like core in a lymphocyte-rich structure. Given the dual nature of ALK1 expression in both blood vessels and TAMs, it is tantalizing to speculate about an active role of this signaling pathway in dampening the anti-tumor potential offered by TLS. In support of this idea, administration of ALK1-Fc significantly upregulated the expression of *Tnfsf14*/LIGHT, an essential chemokine that drives the formation of high endothelial venules (HEVs) and TLS (53). This modulation underlies that HEVs may not respond to conventional pro-angiogenic and -lymphangiogenic signals for their genesis; thus, whether ALK1 is functionally implicated in the development of HEVs remains to be established.

A proportion of mice in our *in vivo* 4T1 setup did progress even in the combined treatment group, reminiscent of the well-known heterogeneity of response to immunotherapy in patients. In this respect, inspection of the longitudinal sc-RNA-seq human melanoma dataset confirms that *ACVRL1* expression follows that of PD-L1 and PD-L2. Interestingly, *Pdcd1lg2*/PD-L2 was consistently upregulated following ALK1 inhibition in the MMTV-PyMT model. The exact role of PD-L2 in respect to PD-L1 is still unclear, with mounting evidence about immune-independent functions of PD-L2 adding to the debate (54). Our experimental data do not seem to support a coordinate expression and utility of these two ligands, as for example SMAD5 is seemingly involved in the transcriptional control of *CD274* only but not *PDCD1LG2*, despite their genomic proximity. From the clinical perspective, this large melanoma cohort also validates that ALK1 expression in TAMs correlates with resistance, with important clinical implications in terms of eligibility of treatment. This observation is particularly relevant in the search for additional companion targets to ALK1 inhibition. For example, *HAVCR2*/TIM-3 is expressed by T-cell to enforce a crucial immune checkpoint function associated with acquired resistance to anti-PD-1 therapy in lung adenocarcinoma (55). Ongoing clinical trials are evaluating the efficacy of agents against TIM-3: while most of the expected effects derive from lymphocytes, these studies should expand to a broader immune landscape, with a thorough characterization of both T cell-dependent and -independent functions (56). In consideration of our *in vivo* results describing a rewired immune landscape, including an increased frequency of T-cells, further studies to assess the role of TIM-3 inhibition in the framework of ALK1 blockade are warranted.

Finally, with the aim of providing an optimal patient stratification for therapeutic gain, it will be paramount to address the concordance between endothelial and myeloid ALK1 expression. Our transplantation-based approach limits us in uncoupling the specific contribution of ALK1 signaling in the endothelial *vs* myeloid compartment. To circumvent this, we devised more controlled strategies to dissect these distinct aspects of ALK1 biology. First, the exploitation of the organ-on-a-chip platform recapitulates the intertwined relationship between tissue vascularization and immune cell trafficking. As the depletion of *Acvrl1* in the endothelial compartment significantly hinders the migration of macrophages, it is likely that modulation of ALK1 remodels the expression of adhesion molecules involved in the multi-step migration of leukocytes. In agreement with this, expression of *Selp* (encoding for P-selecting, necessary for tethering and rolling of leukocytes) is downregulated in liver sinusoidal endothelial cells lacking ALK1 (57). Second, to further dismiss the confounding effect of ALK1 at sites of neo-angiogenesis, the administration of the ALK1 ligand trap in tumor-free hosts exposes the importance of this signaling pathway in the bone marrow. Interestingly, our short-term trial in the 4T1 model corroborates the trends observed in trials with a longer duration but expanded on the role of ALK1 in the mobilization of monocytes, as evinced by the significantly lower relative number of macrophages in tumor-free hosts in the ALK1-Fc cohort compared with the IgG2a control cohort. This regulation transcends the specific requirements dictated by tumor evolution and holds a broader relevance for pathologies that rely on hematopoietic dysfunction for their medical evolution, including the human hemorrhagic telangiectasia that is caused by congenital mutations in *ACVRL1*. An additional readout is the different magnitude of response based on the modality of treatment: in the neoadjuvant regimen, the effects of ALK1 inhibition are *de facto* abolished by primary tumor-derived molecular cues. In support of this, the promotion of an immunosuppressive TME is a preserved feature of growing malignant masses, as our data indicate that progenitor cells extracted from the bone marrow of the MMTV-PyMT model are also more efficient at monocytopoiesis than progenitors from wildtype littermates (Figure S6D).

In the field of TGF-β-related biology, ALK1 has only recently abandoned the role of a pure mediator of endothelial identity and function in favor of a more multifaceted player in different homeostatic processes and diseases, including normal hair maintenance, pulmonary hypertension, and atherosclerosis (58–61). The additional layer of novelty included in our study concerns the differentiation potential of HPCs and the ability of ALK1 to selectively determine the fate of GMPs during tumor evolution. In this context, it remains to be ascertained whether a blunted ALK1 signaling is a priming event that is retained in the differentiation cascade of these early progenitors or if it can be reverted. Finally, the bone represents the most common metastatic site in non-basal breast cancer. Given the peculiar macrophage signature recently uncovered in patients diagnosed with luminal subtypes (62), future efforts will also aim to define the requirement for ALK1 signaling in bone tropism of cancer cells, as well as the conceivable interplay between the hematopoietic and metastatic niches during tumor evolution. As cancer immunotherapy is rapidly incorporating myeloid modulation, a new wave of clinical trials has projected their efforts in re-programming and re-educating macrophage activity through *ex vivo* polarization, adoptive transfer, engineering, and CAR-macrophage approaches (63, 64). In contrast, our work underscores the potential of ALK1 as a target to act upstream of pro-tumorigenic myeloid infiltration in the tumor mass and thereby ameliorate therapeutic response.

In conclusion, our data underpin the clinical value of ALK1 suppression to promote a multifaceted anti-tumor activity. In anticipation of a revamped interest in resuming the development of pharmacological inhibitors against this receptor, our data warrant additional investigation to deconstruct the stromal hierarchy embedded in ALK1 signaling for therapeutic gain.

## Methods

Detailed information on material and methods is provided as supplementary information due to restrictions in word count.

### Mouse work

All mouse work was performed according to the permits M167-15 and 14122-2020 approved by the local ethical committee for animal experimentation.

### Sex as a biological variable

Our study exclusively examined female mice because the disease modeled is mostly relevant in females as no less than 99% of breast cancer diagnoses in women *vs.* men.

## Data and material availability

The RNA-seq data have been deposited with the GEO accession number GSE26091: Link: https://www.ncbi.nlm.nih.gov/geo/query/acc.cgi?acc=GSE260911.

Values for all data points in graphs are reported in the Supporting Data Values file.

## Conflict-of-interest statement

KP and RSP are listed as inventors on a patent related to the ALK1 inhibitor dalantercept.

## Author contributions

Conceptualization: MB, KP. Methodology: MST, JS, PB, EK, TVP, SaL, SoL, GF, VI, SB, MB. Software: JS, PB, ML. Validation: MST, JS, PB, SoL, MB. Formal analysis: MST, JS, PB, EK, TVP, SaL, CO, ML, GBJ, CR, MB, KP. Investigation: MST, JS, PB, EK, EC, TVP, SaL, GF, VI, SB, JP, MB. Resources: RSP, KP. Data curation: MST, JS, PB, EK, MB, KP. Writing – original draft: MB, KP. Writing – review & editing: MST, JS, PB, EK, EC, TVP, SaL, SoL, GF, VI, SB, JP, CO, ML, SRP, GBJ, CR. Visualization: MST, JS, PB, MB. Supervision: MB, KP. Project administration: MB, KP. Funding acquisition: KP.

## Acknowledgements

KP is the Grosskopf Professor of Molecular Medicine at Lund University. This work was supported by grants from the Swedish Research Council, the Swedish Cancer Society, the Knut and Alice Wallenberg foundation, Swedish State Support for Clinical Research through Region Skåne ALF, the Göran Gustafsson foundation, the Mats Paulsson foundations, the Cancera foundation. We thank the Centre for Translational Genomics (Lund University), and Clinical Genomics Lund (SciLifeLab) for sequencing services; Uppsala Multidisciplinary Center for Advanced Computational Sciences (UPPMAX), and the Swedish National Infrastructure for Computing (SNIC) for computing resources. We thank Dr. Marcus Järås for providing human CD14^+^ monocytes; Prof. Kairbaan Hodivala-Dilke for the generous gift of mLEC cells; Christina Möller for technical assistance; Dr. Gottfrid Sjödahl for access to the pathology slide scanner; Anna Ebbesson for access to the laser capture microdissection instrument and training; Prof. Jonas Larsson and Dr. Luis Quintino for reagent provision; Dr. Cristiana Pires, Dr. Trine Ahn Kristiansen, Dr. Charlotta Böiers, Adriana Seira Calderon Moreno, and Lena Tran for scientific discussions; the technical staff at the Lund Stem Cell Centre FACS and Vector cores. Illustrations were created with Biorender.com.

