## Supplemental materials for "An ALK1-governed monocytic lineage shapes an immunosuppressive landscape in breast cancer metastases"

### Supplementary material

#### Methods

##### Cell culture

###### Breast cancer cells

E0771 and 4T1 cells were purchased from American Type Culture Collection (ATCC, VA, USA), and cultured under standard condition (37 °C, 5% $O_2$). E0771 cells were grown in Dulbecco's Modified Eagle Medium (DMEM, 10-013-CV, Corning, AZ, USA) supplemented with 10% fetal bovine serum (FBS, cat. 10500-064, Thermo Fisher Scientific, MA, USA) and 1% Penicillin-Streptomycin (PenStrep, cat. 30-002-CI, Corning, AZ, USA).

4T1 cells were grown in Roswell Park Memorial Institute (RPMI) 1640 (10-040-CV, Corning, AZ, USA) supplemented with 10% FBS and 1% PenStrep.

During culturing, cells were analyzed for mycoplasma infection. Authentication of the cell line identity was confirmed before *in vivo* experiments through CellCheck 19 and IMPACT™ 2 PCR services (IDEXX Bioanalytics, Germany).

For transplantation purposes, cells in exponential growth phase were detached from culture flasks with trypsin-EDTA (25-050-CI, Corning, AZ, USA) and washed with DPBS (D8537, Sigma-Aldrich, MO, USA) to remove dead cells and debris. Cell pellets were resuspended in fresh medium, counted with a Countess automated cell counter (Invitrogen, CA, USA), and washed twice with PBS. Finally, pellets were resuspended at the desired concentration in cold PBS and kept on ice until inoculation.

##### Endothelial cells

Mouse lung endothelial cells (mLECs) were cultured in flasks pre-coated with 0.1% gelatin solution and maintained in complete Endothelial Cell Growth Medium MV2 (C-22022, Merck, Germany).

##### Bone Marrow-derived Macrophages

Total bone marrow cells were extracted from the femur and tibia of 8-12 week-old female FVB mice, and then differentiated to macrophages. Briefly, $3x10^5$ bone marrow cells were cultured in complete RPMI-1640 medium supplemented with 50 ng/mL M-CSF for 7 days, and differentiated macrophages were used for the transendothelial migration.

##### Lentiviral transduction

Short hairpin RNAs (shRNAs) targeting *Acvrl1* were ordered from Dharmacon (CO, USA) in form of SMARTvector Lentiviral shRNA expressing the GFP reporter gene. We used two shRNAs targeting Acvrl1 (shA07 and shA09) and one non-targeting shControl (shCtrl). Lentiviral particles were generated at the Lund University vector unit facility with high titers and efficiency. Prior to transduction, $1x10^5$ mLEC cells were seeded in a 6-well plate and each well was treated with shCtrl, shA07 or shA09 lentiviral particles and incubated at 37 °C. One-week post-transduction, cells were checked under microscope to check the GFP expression and samples were collected for quantifying $GFP^+$ cells and knock-down efficiency by qPCR.

##### Organo-plate

For the Extra Cellular Matrix (ECM) preparation, rat tail collagen type I (5 mg/mL, 3447-020-01, R&D Systems, MN, USA) was mixed at a ratio of 8:1:1 with 1 M HEPES (25-060-CI, Corning, AZ, USA) and 37 g/L NaHCO3 (S5761-500G, Sigma-Aldrich, MO, USA) producing a 4 mg/mL collagen type I solution. The mixture was prepared on ice and used within less than 10 minutes after preparation. 50 µL of 1X Hanks' Balanced Salt Solution (HBSS) was added to the observation window of a 3-Lane OrganoPlate (4004-400-B, Mimetas, the Netherlands) to prevent chips from drying out. 2.2 µl of ECM gel was added to the middle channel of the chip via the gel inlet followed by addition of 30 µl HBSS to the gel inlet to prevent ECM dehydration. Then, the plate was placed in a humidified incubator for 15 minutes to allow polymerization of the gel. After incubation, HBSS was aspirated from the gel inlets and MLEC cells were seeded into each top channel by addition of 2 µl cell suspension to the top medium inlet. Media was added to the top medium inlet and the plate incubated for 2 hours on its side at a 75º angle to allow cells to attach to the ECM. Media was then added to the top medium outlet and perfusion begun (8 minutes interval, 7º angle) on the Mimetas OrganoFlow® plate rocker (Mimetas, The Netherlands). The plate was incubated under perfusion for 8 days to enable optimal formation of endothelial tube with optimal barrier function of the vessels. Media from the top medium inlet and outlet was changed every other day. Imaging was performed with Laser Scanner Microscope (LSM 710, Zeiss, Germany) using the ZEN Black Software to take Z-stacks with a 10x objective. All Z-stacks were reassembled in a 3D reconstruction using the FIJI (65) plugin.

##### Barrier integrity assay

After 8 days of incubation, the integrity of the EC tubing was tested. Fluorescent working solution was prepared by diluting 25 ng/mL TRITC-dextran 4.4 kDa (T1037, Sigma-Aldrich, MO, USA) stock solution and 25 ng/mL FITC-dextran 150 kDa (46946, Sigma-Aldrich, MO, USA) stock solution with mLEC culture media so that the concentration of each fluorescent compound was 0.5 mg/mL in the final working solution. A 5-minute "wetting" step was performed by adding 50 µL of mLEC media to perfuse all the inlets and outlets. mLEC culture media was then aspirated and 20 µL of mLEC culture media was added to the gel inlet, gel outlet, bottom medium inlet and bottom medium outlet. 40 µl and 30 µl of the final fluorescent working solution were added to the top medium inlet and outlet, respectively. Dye retention was imaged on a ZEISS LSM 710 with a 10X objective. Barrier function was quantified by measuring the fluorescence in each channel in FIJI and then calculating the ratio of fluorescence in the vessel channel (top channel) versus fluorescence in the ECM channel (middle channel).

#### Mouse work

##### Therapeutic regimens

Treatment schedule, dosage, and delivery of RAP-041 (Acceleron Pharma, MA, USA, now Merck) were established in previous work (9), with a twice weekly intraperitoneal (i.p.) injection of 12 mg/kg body weight of RAP-041. InVivoMAb mouse IgG2a isotype control (BE0085, BioXCell, NH, USA) was administered with the same concentration and regimen as RAP-041. InVivoPlus anti-mouse PD-1 and anti-mouse CTLA-4 (BP0146 and BP0164, BioXCell, NH, USA) were supplied once weekly via i.p. injection at 10 mg/kg body weight. All compounds were diluted in PBS, and further delivered in different combinations as highlighted in the study design for each preclinical trial.

##### Therapeutic trials

*E0771 adjuvant*: $5x10^5$ E0771 cells/100 µl of PBS were transplanted in the 4th mammary fat pad of 8-week-old C57BL/6 female mice (Harlan, IN, USA, or Janvier Labs, France). Approximately 3 weeks after, primary tumors reached 13 mm in the longest diameter, a measurement that was previously shown to induce the outgrowth of lung metastases (66). At this stage, pre-operative care included shaving, application of eye drops (Viscotears 2mg/g, Laboratoires Thea, France) and administration of painkiller (Norocarp, 5 mg/kg body weight). During the whole procedure, mice were kept under general anesthesia with isoflurane (002185, Zoetis, NJ, USA). Primary tumors (including the mammary fat pad whenever possible) were excised through a 5-mm surgical incision in the abdominal cavity. If the tumor tissue grew close connection to internal organs, a cauterization pen (AA01, Bovie Medical Corporation, FL, USA) was used to prevent bleeding. Following the cleaning of the surgical area with chlorhexidine acetate (0,5 mg/ml, Fresenius Kabi, Germany), wounds were closed with Vetbond Tissue Adhesive (1469SB, 3M, MN, USA) or sutures (Coated VICRYL 6-0, J384H, Ethicon, NJ, USA), and mice monitored for recovery. Post-operative Norocarp was administered once a day for up to 3 days after the surgery. Five to 7 days after surgery, mice were randomized into two groups receiving either IgG2a or RAP-041 for up to 4 weeks, with the same treatment schedule described before. Necropsy to assess the lung metastatic burden was performed also on mice that did not reach the experimental end-point.

*4T1 adjuvant*: 5x$10^4$ 4T1 cells/50 μl of PBS were orthotopically transplanted in 8-week-old BALB/c female mice (The Jackson Laboratory, ME, USA). Ten to 12 days after inoculation, primary tumors reached an average of 6 mm in diameter, thereby classifying eligible mice for surgical resection and relative post-operative care as described above. At the end of a recovery period of 5 days, mice were randomized into four groups receiving either IgG2a, RAP-041, dual anti-PD-1 + anti-CTLA-4 blockade, or RAP-041 + anti-PD-1 + anti-CTLA-4 for 3 weeks. Necropsy to assess the lung metastatic burden was performed also on mice that did not reach the experimental end-point.

*4T1 adjuvant early resection*: $10^5$ 4T1 cells/50 μl of PBS were orthotopically transplanted in 9-week-old BALB/c female mice. Seven days after the inoculation, tumors reached an average diameter of 4.5 mm, and were resected as described previously. Five days after surgery, mice were randomized into two groups receiving either IgG2a or RAP-041 for 3 weeks.

*4T1 short-term*: 5x$10^4$ 4T1 cells/50 μl of PBS were orthotopically transplanted in 8-week-old BALB/c female mice. Twelve days after inoculation, primary tumors reached an average of 6 mm in diameter, thereby classifying eligible mice for surgical resection and relative post-operative care as described above. After 3 days of recovery, mice were randomized into two groups receiving either IgG2a or RAP-041. Compounds were administered three times during the course of 7 days, and mice sacrificed 24 hours after the last treatment. In the neoadjuvant setup, treatment commenced when tumors reached approximately 100 $mm^3$ in volume.

*MMTV-PyMT neoadjuvant*: Eleven-week-old MMTV-PyMT female mice were randomized to receive IgG2a, PD-1 or RAP-041 + anti-PD-1 according to the regimen described above. Mice were treated for four weeks until sacrifice at 15 weeks of age. Tumors were measured before randomization to ensure comparable sizes among groups, and further recorded once weekly with an electronic caliper.

*E0771 neoadjuvant*: $5x10^5$ E0771 cells/100 µl of PBS were orthotopically transplanted in 8-week-old C57BL/6 female mice. When tumors became palpable (approximately 40 $mm^3$), mice were randomized to receive IgG2a control, RAP-041, anti-PD-1, anti-CTLA-4 or a combination of RAP-041 with either anti-PD-1 or anti-CTLA-4, with the treatment dosage and schedule described before. Compounds were administered for 2 weeks, corresponding to the humane end-point in the ethical permit (tumor volume = 1 $cm^3$).

#### Tissue harvesting and processing

For MMTV-PyMT and E0771 models, the circulatory system was flushed with 10 ml PBS through ventricular injection. For 4T1-based experiments, mice were not perfused with PBS to allow for best comparison with primary tumors at the time of resection. Lungs, tumors, and bones from the hind legs were excised and kept in cold PBS + 2% fetal calf serum (FCS) until processed. Peripheral blood was collected *via* cheek-bleed in EDTA-coated tubes (20.1339.100, Sarstedt, Germany) at specific time points as indicated in the study design in the corresponding figures, as well as at the experimental endpoint right before cervical dislocation. Upon necropsy, organs were collected for downstream preparations. For RNA work, a portion of the tissue was excised and directly placed in liquid $N_2$. For frozen sections, tissues were stored in 30% v/v sucrose in PBS for 24 hours, followed by embedding in optimal cutting temperature (OCT) compound (45830, Histolab, Sweden). For histology and immunohistochemistry, tissue was placed in zinc formalin fixative (Sigma-Aldrich, MO, USA) for 24 hours. Samples were stored in 70% ethanol until paraffinization with a benchtop carousel tissue processor (Microm STP120, Thermo Fisher Scientific, MA, USA), followed by the generation of formalin-fixed paraffin embedded (FFPE) blocks with a tissue embedding console system (Tissue-Tek, Sakura Finetek, Japan).

#### Tissue staining and analysis

##### Hematoxylin and Eosin staining (H&E)

5 μm-thick FFPE sections were rehydrated in xylene and subsequently exposed to absolute ethanol, 96% ethanol, 70% ethanol, and water. Samples were stained with Harris hematoxylin for 5 minutes at RT, followed by washes with tap water, and finally exposed to eosin for 1 minute at RT. Tissue sections were dehydrated with a reverse protocol from 70% ethanol to xylene, prior to mounting with Cytoseal 60 (Thermo Fisher Scientific, MA, USA). Full-tissue images were acquired with a NanoZoomer S60 digital slide scanner, followed by visualization for annotation and analysis with the NDP.view2 software (Hamamatsu Photonics, Japan).

##### Immunohistochemistry (IHC)

5 μm-thick FFPE sections were rehydrated in xylene and subsequently exposed to absolute ethanol, 96% ethanol, 70% ethanol, and water. Antigen retrieval was performed with a 2100 retriever (Prestige Medical, UK) in appropriate pH6/pH9 buffer (AR600250ML and AR900250ML, both from Akoya Biosciences, MA, USA). Endogenous peroxidase activity was quenched with BLOXALL (SP-6000, Vector Laboratories, CA, USA), 10 minutes at RT, followed by incubation with CAS block (008120, Life Technologies, CA, USA), 1 hour at RT in a humidified chamber. Primary antibody diluted in CAS block was applied for 1 hour at RT in a humidified chamber. Sections were washed with PBS-Tween20 (PBST) and incubated with a mouse- or rabbit-specific HRP-tagged SignalStain Boost IHC detection reagent (8114S and 8125S, Cell Signaling Technology, CA, USA) for 30 minutes at RT in a humidified chamber.

Tissue sections were washed with PBST before supplying diaminobenzidine (DAB, SK-4105, Vector Laboratories, CA, USA) for the chromogenic development of the labeling. Following counterstaining with Harris hematoxylin, tissue sections were dehydrated to xylene and mounted with Cytoseal 60. Full-tissue images were acquired with a NanoZoomer S60 digital slide scanner. Primary antibodies: anti-CD3 1:150 (ab16669, Abcam, UK), anti-CD45 1:100 (553076, BD Pharmingen, CA, USA).

##### Multiplexed immunohistochemistry (mIHC)

A customized 6-plex antibody panel (Table S8) was developed to concomitantly detect blood vessels and four major immune cell types in murine tumor tissue.

Briefly, 5 µm-thick FFPE sections were placed at 60 °C, O/N, to melt the excess of paraffin, and further rehydrated by subsequent washes in xylene, absolute ethanol, 96% ethanol, and water. Sections were fixed in neutral buffered formalin (NBF, HT501128-4L, Sigma-Aldrich, MO, USA), 20 minutes, RT, before heat-induced antigen retrieval (HIER) at a pH specific for the first antibody of the panel. Each staining cycle is composed of a blocking step, followed by incubation with a primary antibody in blocking buffer. After a series of washes in PBS-Tween20, a species-specific HRP-conjugated secondary antibody was applied. Then, sections were labeled with an OPAL fluorophore-conjugated tyramide substrate (all from Akoya Biosciences, MA, USA). Samples were washed in PBS-T, and a HIER-based stripping was performed to quench any residual peroxidase activity, as well as to prepare the tissue for the next staining cycle. At the end of six cycles, sections were counterstained with spectral DAPI, and mounted with ProLong Diamond antifade mountant (P36961, Thermo Fisher Scientific, MA, USA). Slides were acquired with a PhenoImager HT instrument and analyzed with the built-in software (PhenoChart, InForm 2.6), and the phenoptrReports R package (all from Akoya Biosciences, MA, USA).

##### Dual in situ hybridation (ISH) combined with immunofluorescence

A protocol for the concomitant detection of *ACVRL1* mRNA, as well as CD31 and CD45 protein markers on human breast cancer tissue was developed ad hoc. The procedure was based on the RNAscope Multiplex Fluorescent V2 assay combined with immunofluorescence – Integrated co-detection workflow (323180, 322381, 323100, 322809 and 310091, all from ACD Advanced Cell Diagnostic, CA, USA). Briefly, 5 µm-thick FFPE sections of human breast cancer tissue underwent pretreatment, blocking and target retrieval. A cocktail of CD31 and CD45 antibodies (3528S and 13917S, Cell Signaling Technology, CA, USA) was added to the sections, and incubated O/N at 4 °C in a humidified chamber. Protease treatment preceded the actual RNA-ISH assay with a ready-to-use RNAscope Hs-*ACVRL1* probe as well as positive and negative control probes (559221, 320861 and 320871, ACD Advanced Cell Diagnostic, CA, USA). Next, tyramide-based detection of the two protein markers was performed. The primary antibody was recognized by a species-specific HRP-conjugated secondary antibody (Cell Signaling Technology, CA, USA), followed by an OPAL fluorophore (FP1487001KT, FP1495001KT and FP1497001KT, Akoya Biosciences, MA, USA). Tissue sections were then exposed to a 2% peroxide quenching solution for 15 minutes at RT to block any residual HRP activity from one antibody detection cycle to the other. The same labeling procedure was performed for the second protein marker. Tissue sections were mounted with ProLong Diamond antifade mountant. Finally, slides were acquired with a PhenoImager HT instrument, and analyzed with complementary PhenoChart and InForm 2.6 software.

##### Flow cytometry and FACS sorting

For cell surface staining, cells were collected, washed and prepared as single cell suspension. Prior to analysis or cell sorting cells were labeled with combination of the fluorochrome-conjugated anti-mouse antibodies as stated in the below table for 45 minutes at 4°C and washed afterwards using cold FACS buffer (PBS + 5% FBS). All data were collected with fluorescenceactivated cell sorter (FACS) X20 or LSRII analyzer (Becton Dickinson, NJ, USA). Cell sorting was performed using AriaIII or Melody (Becton Dickinson, NJ, USA) and all the analysis was done with the FlowJo software (OR, USA). For the immune cell sort, cells were directly sorted into RLT buffer prepared for RNA extraction.

| Antibody | Conjugate | Clone | Source |
|---|---|---|---|
| CD45 | Alexa700 | 30-F11 | BioLegend |
| CD19 | pacific blue | 6D5 | BioLegend |
| CD8 | FITC | 53-6.7 | BioLegend |
| CD335-NKP46 | BV421 | 29A1.4 | BioLegend |
| B220 | PE-cy7 | RA3-6B2 | BioLegend |
| CD11c | FITC | N418 | BioLegend |
| MHC Class II (I-A/I-E) | PE/Cy7 | M5/114.15.2 | BioLegend |
| CD64 | APC | X54-5/7.1 | BioLegend |
| Ly-6C | BV421 | HK1.4 | BioLegend |
| Ly-6G | BV785 | 1A8 | BioLegend |
| CD4 | BV786 | GK1.5 | BD |
| CD11b | BV605 | M1/70 | BD |
| CD3e | BV711 | 145-2C11 | BD |
| CD117(c-Kit) | APC-cy7 | 2B8 | BioLegend |
| Ly-6A/E (Sca1) | BV421 | D7 | BioLegend |
| CD150 (SLAM) | BV605 | TC15-12F12.2 | BioLegend |
| CD41 | FITC | MWReg30 | BD Pharmingen |
| CD16/32 (FcGR) | APC | 93 | BioLegend |
| CD48 | PE/Cy7 | HM48-1 | BioLegend |
| Ly-6G/Ly-6C (Gr-1) | PE/Cy5 | RB6-8C5 | BioLegend |
| CD45R/B220 | PE/Cy5 | RA3-6B2 | BioLegend |
| CD3e | PE/Cy5 | 145-2C11 | BioLegend |
| NK1.1 | PE/Cy5 | PK136 | BioLegend |

#### Bone marrow c-Kit enrichment and CFC assay

Colony-forming cell (CFC) assay was performed by plating magnetically enriched BM-cKit$^+$ cells using mouse CD117 microbeads (130-097-146, Miltenyi Biotec, Germany). Briefly, BM cells were resuspended in 100 µl medium per 100x10$^6$ cells and 2.5 µl beads per 100x10$^6$ cells was added to each sample. After 20 minutes incubation on ice, and washing steps, c-Kit$^+$ cells were run through LS columns (130-042-40, Miltenyi Biotec, Germany) designed for positive selection of magnetically labelled cells. Enriched c-Kit$^{+}$ cells were then plated on Methylcellulose-based medium with recombinant cytokines for mouse cells (MethoCultTMGF M3434, STEMCELL technologies, BC, Canada) in 35 mm culture dish at 37 °C, 5% $CO_2$. Colony forming units were scored at 14 days in culture. For hematopoietic stem and progenitor cell staining, total BM cells were stained with HSC (LSK-SLAM) cell surface markers (Lineage(B220, NK1.1, Gr1 and CD3e), cKit, Sca1, CD48, CD150, CD16/32 and CD41 for 45 minutes at 4°C and washed afterwards using cold FACS buffer (PBS + 5% FBS). 7AAD (BD Pharmingen #51-68981E) was added to individual samples prior analysis in order to exclude the dead cells.

#### RNA work

##### RNA extraction from FFPE tissue for PCR array

Freshly cut FFPE tissue section from preclinical trials with ALK1-Fc (13) were cut to obtain approximately 40 µm of specimen. RNA was extracted from this starting material with the RNeasy FFPE kit (73504, Qiagen, Germany) according to the manufacturer's datasheet. Eluted RNA was used as input for the RT2 profiler 96-well PCR array for T-cell and B-cell activation (PAMM-053, Qiagen, Germany) according to the manufacturer's datasheet.

##### Manual dissection and RNA isolation

Lung macrometastases were manually dissected on ice with a sterile surgical blade. Approximately 30 mg of metastatic tissue was placed in a 2 ml tube with 600 µl of RLT buffer, 60 µl β-mercaptoethanol, and 3 µl anti-foam DX reagent (Qiagen, Germany). A 5-mm steel bead (Qiagen, Germany) was used to fully homogenize samples with a TissueLyser LT (Qiagen, Germany), 50 oscillations/s for 3 minutes. Lysate was passed through a QIAshredder (Qiagen, Germany), and centrifuged at 16000 rcf for 1 minute. The flow-through was used for the RNA extraction with a RNeasy mini kit (Qiagen, Germany), according to the standard protocol provided by the manufacturer.

#### Laser-capture microdissection (LCM) and RNA isolation

Snap-frozen metastases-bearing lungs from the E0771 adjuvant trial were sectioned at a thickness of 16-18 μm and let dry on polyethylene naphthalene (PEN) membrane 1.0 slides (Zeiss, Germany). Prior to their use, membrane slides were dry-heated for 4 hours at 180°C to ensure RNase-free conditions. Moreover, membrane slides were exposed to ultraviolet (UV) light treatment at 254 nm for 30 minutes to improve the adhesion of the specimen. Tissue sections were fixed in 70% ethanol for 3 minutes at RT, followed by staining with 1% cresyl violet in 50% ethanol for 30 seconds. The excess of dye was removed, and slides were sequentially dipped in 70% and absolute ethanol before a final air-drying step. For long term preservation, stained slides were stored at -80 °C in air-tight containers. To avoid condensation, lids were opened only after specimens fully thawed.

Metastases were captured with a PALM microdissection system (Zeiss, Germany) on a 500 μl tube with adhesive cap, with a total dissected area per sample ranging 4-9 $mm^2$. Samples were immediately processed with a modified protocol based on the MinElute PCR purification kit (Qiagen, Germany). In the first step, samples were diluted in 350 μl of RLT buffer, and further incubated upside down for 30 minutes at RT. Samples were spun down at 16000 rcf for 5 minutes and transferred to a 1.5 ml tube to complete the extraction with a RNeasy micro kit (Qiagen, Germany), according to the standard protocol provided by the manufacturer.

#### RNA quality controls

Following the RNA extraction, 1 μl of the elute was used for quantification of the total RNA yield, as well as 260/280 and 260/230 ratios, with a Nanodrop 2000 spectrophotometer (Thermo Fisher Scientific, MA, USA). For samples with concentration below 40 ng/μl, an additional aliquot was used to determine the RNA concentration with a Qubit fluorometer (Thermo Fisher Scientific, MA, USA). Before sample selection, evaluation of RNA degradation and their relative RNA integrity number (RIN) was performed with a 2100

Bioanalyzer system equipped with eukaryote total RNA Nano or Pico assay chips (Agilent Technologies, CA, USA). Total RNA was stored at -80 and delivered to the Center for Translational Genomics (CTG, Lund, Sweden) for library preparation, sequencing and data pre-processing.

##### RNA-seq data sequencing and processing

RNA-seq data sequencing and processing was performed by CTG. Briefly, the raw data was generated using NextSeq 500 (SY-415-1001, Illumina Inc. CA, USA) and processed using Illumina Dragen RNA pipeline for RNA quantification on transcript and gene level, as per standard protocol (Illumina Inc., CA, USA). We used tximport (v1.23.4) to import and summarize transcript-level abundance estimates for gene-level downstream analysis. We demultiplexed the raw data to FASTQ files using bcl2fastq (v2.20.0 Illumina, RRID: SCR_015058) followed by a quality assessment of the FASTQ files using FastQC[6]. The reads were aligned to the mouse reference genome GRCm38. Reference genome and annotation (GTF file) were downloaded from the Ensemble database release 99 (67). We used default settings for all tools, unless otherwise specified.

##### RNA-seq differential expression analyses

We used Deseq2 (68) (v1.26) to test for differential expression between treatments. We estimated the dispersion parameters using all samples, but differential gene expression was tested using a simple design doing multiple pairwise comparisons. We used the results function with the contrast argument followed by the LfcShrink function suing type=”normal”.

##### Gene Set Enrichment Analysis

For the murine pipeline, we used the MsigDB R package (v7.2.1), that provides Molecular Signatures Database (MsigDB) gene sets (69), in combination with the fast gene set enrichment analysis (fGSEA) package (v1.12.0) which implements an algorithm to perform fast preranked gene set enrichment analysis (70). We ranked all genes by log fold change. fGSEA enrichment analysis was performed for 6 different gene set collections, namely Hallmark pathways, oncogenic signatures (C6), computational (C4), GO (C5), immune (C7) and position (C1) gene sets.

The overrepresentation analysis was performed using goseq (v1.38.0) using the annotation from the hg38 from the TxDb.Hsapiens.UCSC.hg38.knownGene package (71) (V3.10). All significantly differential expressed genes were included in the analyses (alpha <0.05).

For TCGA data, a ranked list and relative RNK file base don co-expression with *ACVRL1* were obtained by enquiring the "cBioPortal for cancer genomics" (24, 25) in the Invasive Breast Carcinoma (Firehose Legacy study) cohort of The Cancer Genome Atlas (TCGA) repository in April 2022. In this case, gene ranking was based on Spearmans's R coefficient. The RNK list was used as input for gene set enrichment analysis (GSEA) preranked analysis Hallmarks pathways.

##### scRNA-Seq Datasets

All scRNA-seq data used in this study were publicly available. The inDrop breast cancer scRNA-seq dataset (32) was obtained from the Gene Expression Omnibus (GEO) repository with the accession codes GSE114725. The Smart-Seq2 melanoma scRNA-seq dataset (42) was downloaded from the TISCH2 database (72). The 10x Genomics Chromium breast cancer scRNA-seq dataset (29) of all cell types and the reclustered myeloid cells from this dataset subset were downloaded from the Single Cell Portal – Broad Institute. The 10x Genomics Chromium breast cancer TNBC and $ER^+$ scRNA-seq datasets (31) were downloaded from figshare (https://figshare.com/articles/dataset/Data_R_code_and_output_Seurat_Objects_for_single_cell_RNA-seq_analysis_of_human_breast_tissues/17058077). The 10x Genomics Chromium Human Breast Cell Atlas (HBCA) myeloid cell scRNA-seq dataset (27) was downloaded from CELLxGENE (https://cellxgene.cziscience.com/collections/4195ab4c-20bd-4cd3-8b3d-

65601277e731). The CITE-seq human bone marrow dataset (28) was installed using the SeuratData (73) package (version 0.2.2).

##### scRNA-seq Data Preprocessing

For the analysis of the inDrop breast cancer dataset comprising FACS-sorted $CD45^+$ cells, we removed genes that had zero counts in all cells and further filtered the count matrix by only keeping genes, which were expressed by 10 or more cells. Then, we followed the preprocessing steps of Cheng *et al.* (30), but retained only cells that comprised the five macrophage cell clusters (Macro_C1QC, Macro_CX3CR1, Macro_ISG15, Macro_SPP1, and Macro_VCAN), which had been identified upon their reanalysis of the Azizi and colleague's dataset (29). The bone marrow dataset was preprocessed exactly as in the Weighted Nearest Neighborhood (WNN) vignette using the Seurat package (73) (version 4.4.0) to create an UMAP using the weighted combination of the RNA and antibody-derived tags assays. Processed expression matrices, UMAP coordinates, existing cell annotations, and clinical metadata regarding, *e.g.* treatment regimens, for all other datasets were used as specified in the original publication sources. We reassessed some quality control metrics, *e.g.* percentage of mitochondrial reads, to ensure that gene- and cell-level filtering had been performed appropriately.

##### scRNA-seq Data Analysis and Visualization

All datasets were imported and further analyzed using Seurat. Gene expression levels, t-SNE and UMAP representation of cell clusters, and AddModuleScores (a function in Seurat to calculate gene set module scores) were visualized using the scCustomize (version 0.7.0), the SCpubr (version 2.0.2), the dittoSeq (version 1.9.0), and the ggplot2 (version 3.4.4) packages. The average expression of genes was calculated using the AverageExpression function in Seurat.

To construct a robust signature to accurately represent *ACVRL1*-positive TAMs, a strategy based on differential expression using two datasets were employed. Seurat's FindMarkers function (non-parametric Wilcoxon rank sum test) was used to find differentially expressed genes (n= 400, adjusted p-value < 0.05) between *ACVRL1*-positive (expression > 0.0) and *ACVRL1*-negative (expression <= 0.0) cells in the macrophage cell populations from Azizi *et al*. (29) (Table S4). The ACVRL1-positive DEG list was further filtered to only include genes (n = 206) with a log2FC > 0.3 and over 0.2 in difference in the fraction of detection (min.pct.diff = 0.2) between the *ACVRL1*-positive and *ACVRL1*-negative groups. Then, the FindAllMarkers function (non-parametric Wilcoxon rank sum test) was used to find DEGs between all cell types in the TNBC dataset (31), and the macrophage-specific genes (n = 31, adjusted p-value < 0.05, log2FC > 2.0, min.pct.diff = 0.3, as compared to cancer cells, fibroblasts/pericytes, endothelial cells, B-cells, T/NK-cells, and plasma cells; Table S5) were compared to the *ACVRL1*-positive list from Azizi *et al*. (29), resulting in four genes (*APOC1*, *FCER1G*, *MMP9*, and *SPP1*) that met all these filtering criteria.

##### scRNA-Seq Data Enrichment Analysis

Macrophage meta-programs from The Curated Cancer Cell Atlas (33) (3CA) were downloaded from the 3CA webpage and used in the overrepresentation analysis. The Genekitr package (version 1.2.5) was used to calculate enrichment scores for the meta-programs in the gene list (n= 400, adjusted p-value < 0.05) associated with the *ACVRL1*-positive TAM population (Table S6).

##### ChIP-Seq Data Analysis

ChIP-Seq bigwig-files of the SMAD5, P300, and CTCF transcription factors and bigwig-files of histone marks (H3K27ac, H3K9ac, H3K4me3, and H3K36me3) from analysis of a human lymphoblastoid cell line (ENCODE biosample ID: GM12878) were downloaded from the ENCODE (44) project website. Peak profiles of histone modifications and the transcription factors were visualized using karyoploteR (version 1.20.3) on gene regions of interest using the TxDb.Hsapiens.UCSC.hg19.knownGene (version 3.2.2) package.

#### Prognostic Analysis

Breast cancer data gene expression profiles and matched clinical information from the TCGA and the METABRIC consortiums were obtained from the cBioPortal database. The patients were divided into two risk groups based on the median value of the mean expression of the *ACVRL1* signature, which had been divided by the expression of *PTPRC* to account for the immune infiltration of each sample. The survival analysis was performed using the Surv and survfit functions in the survival package (version 3.5-5). The Kaplan–Meier curves were drawn using the ggsurvplot function in survminer (version 0.4.9). The log-rank test was used for statistical assessment of the survival time analyses. We also applied a Cox proportional hazard model to correct for key breast cancer clinical parameters, *i.e.* age, stage, and estrogen receptor status. Summary tables of the Cox proportional hazard model was produced using the gtsummary package (version 1.7.2).

### Figure S1. Related to Figure 1

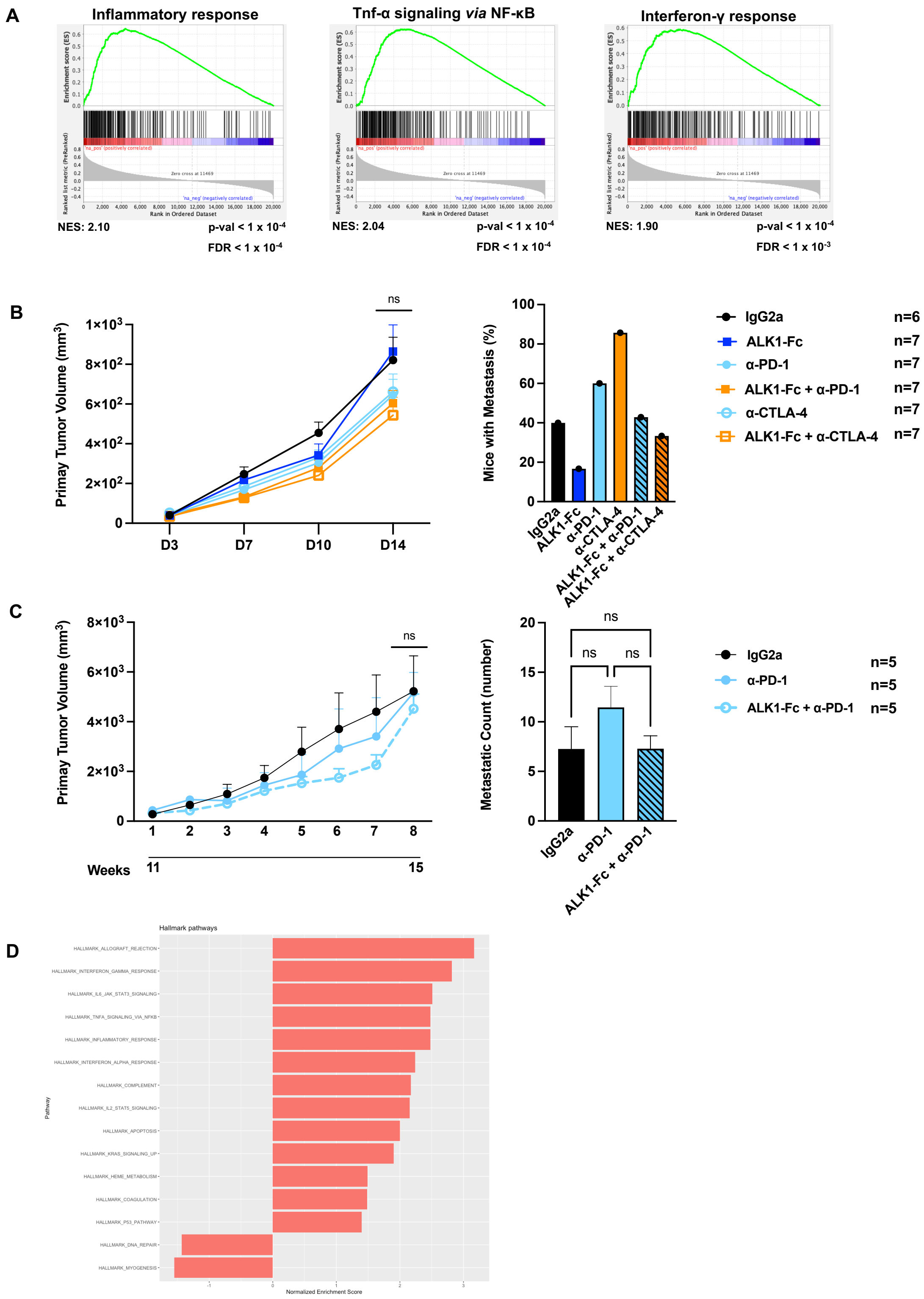

Figure S1.

(A): Representative enrichment plots for Inflammatory response (left), TNF-α signaling via NF-κB (middle), and Interferon-γ response (right) from the GSEA analysis of genes co-expressed with *ACVRL1* in the Invasive Breast Carcinoma cohort of The Cancer Genome Atlas. NES: normalized enrichment score; FDR: false discovery rate. (B): Primary tumors were established by orthotopically transplanting E0771 cells in syngeneic C57BL/6 mice. Neoadjuvant administration of ALK1-Fc, anti-PD-1, anti-CTLA-4, or a combination of ALK1-Fc with either immunotherapeutic agent commenced when tumors were palpable. Primary tumor volume curves (left), and proportion of mice with lung metastasis at the experimental end point (right). Data displayed as mean with SEM. P-value: ANOVA test (ns: not significant). (C): 11-week-old transgenic MMTV-PyMT mice were randomized to receive neoadjuvant ALK1-Fc, anti-PD-1, or ALK1-Fc + anti-PD-1 for four weeks. Primary tumor volume curves (left), and average number of lung metastases at the experimental end point (right). Data displayed as mean with SEM. p-value: ANOVA test (ns: not significant). (D): GSEA hallmark analysis of RNA from laser capture microdissected lung metastases from the adjuvant E0771 trial, ALK1-Fc *vs* IgG2a.

#### Figure S2. Related to Figure 2

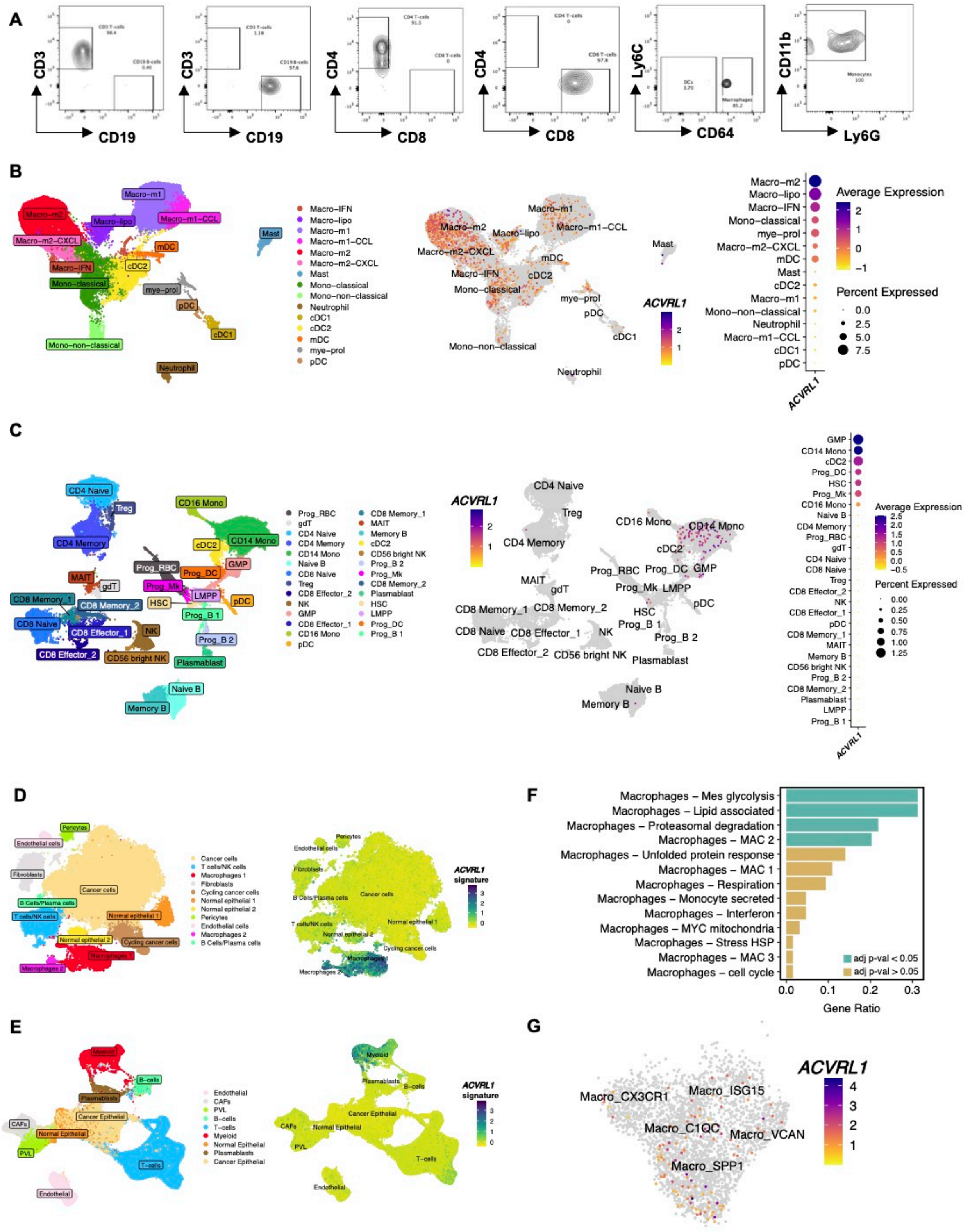

Figure S2.

(A): Purity check of the isolation of immune populations through FACS-sorting of primary MMTV-PyMT tumors dissociated to single cell suspensions. (B): Expression of *ACVRL1* in annotated myeloid cells from the human breast atlas (27). The expression of *ACVRL1* was imposed on the UMAP with annotations and presented as a dot-plot with averaged expression within each cluster. (C): Expression of *ACVRL1* in the CITE-seq dataset of the healthy human bone marrow (28). The expression of *ACVRL1* was imposed on the WNN UMAP with annotations, and further presented as a dot-plot with averaged expression within each cluster.

(D-E): Overlay of a 5-gene *ACVRL1* signature specific for TAMs onto all cell types of two human breast cancer scRNA-seq datasets (31, 32). (F): Gene ontology for metaprograms of the 400 significantly DEGs between $ACVRL1^{+}$ *vs* $ACVRL1^{-}$ TAMs from the breast-specific immune phenotype atlas (29). (G): Feature plot of *ACVRL1* expression imposed on the UMAP of the scRNA-seq atlas of immune phenotypes (29, 30).

### Figure S3. Related to Figure 3

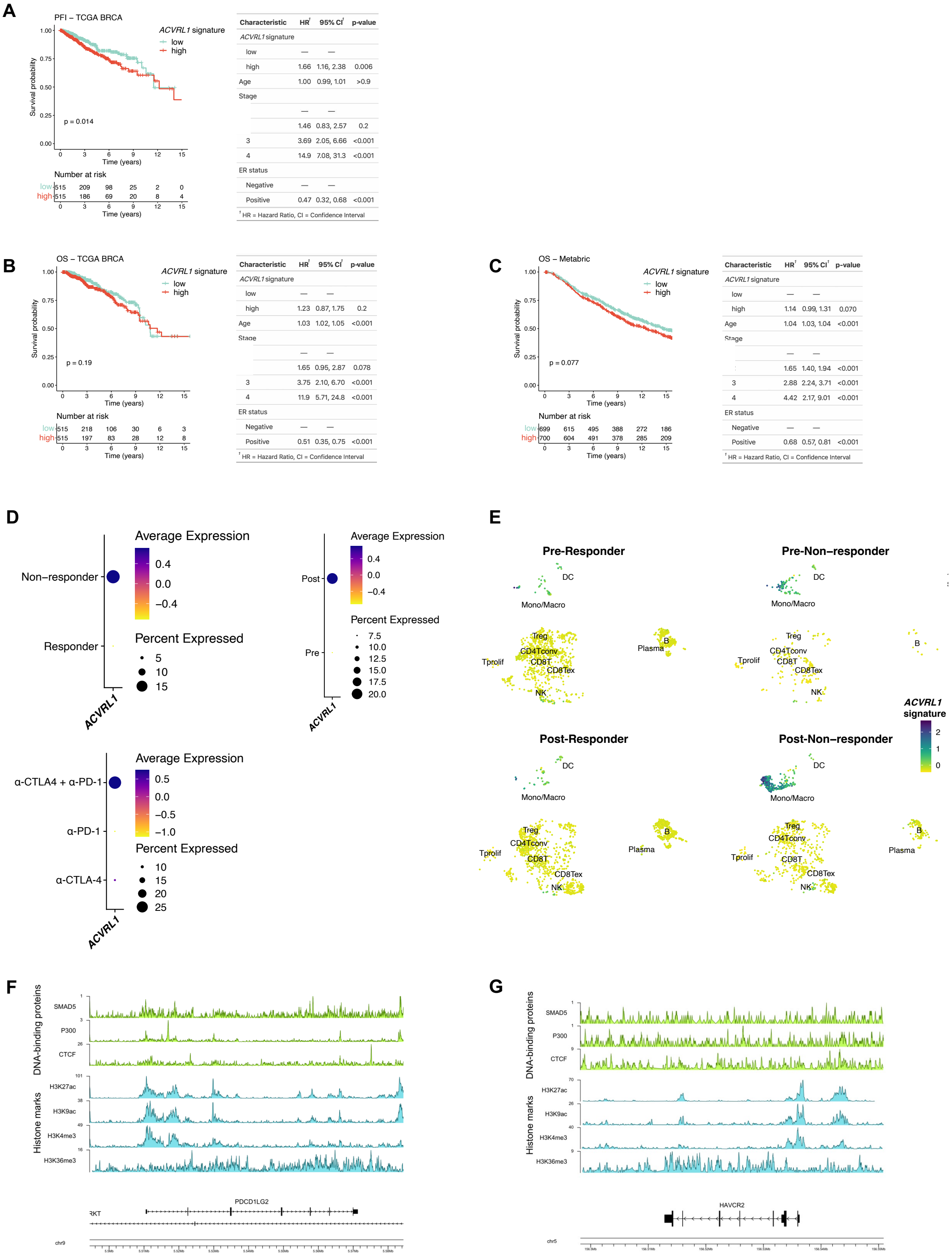

Figure S3.

(A-C): Survival analysis in the TCGA BRCA (39) (A) and METABRIC (40) (B) datasets. Patients were stratified into two risk groups based on the median value of the mean expression of a TAM-specific *ACVRL1* signature. The Kaplan-Meier curves show the progression-free interval (PFI; A), and overall survival (OS; B-C) probabilities of the high (red) and low (green) signature expression groups in the two cohorts. p-value: log-rank test. The tables summarize the relative Cox proportional hazard model analysis for each cohort. (D-E): Expression of *ACVRL1* in a $CD45^+$-restricted scRNA-seq compendium of 48 melanoma patients treated with immune checkpoint inhibitors (42). The average expression of *ACVRL1* is presented in dot-plots based on time point, response, and treatment arm (D). The average expression of the 5-gene signature in the combined CTLA-4 and PD-1 inhibition was imposed on the UMAP, and further displayed as a feature plot based on the response to treatment (responder *vs* non-responder) before (pre) and after (post) treatment, thereby creating four different groups (E). (F-G): ChIP-seq plot from human hematopoietic tissue (43, 44). Visualization of DNA-binding protein and histone marks in the promoter region of *PDCD1LG2* (F), and *HAVCR2* (G). SMAD5 activity is the direct downstream modulator of canonical ALK1 signaling.

### Figure S4. Related to Figure 4

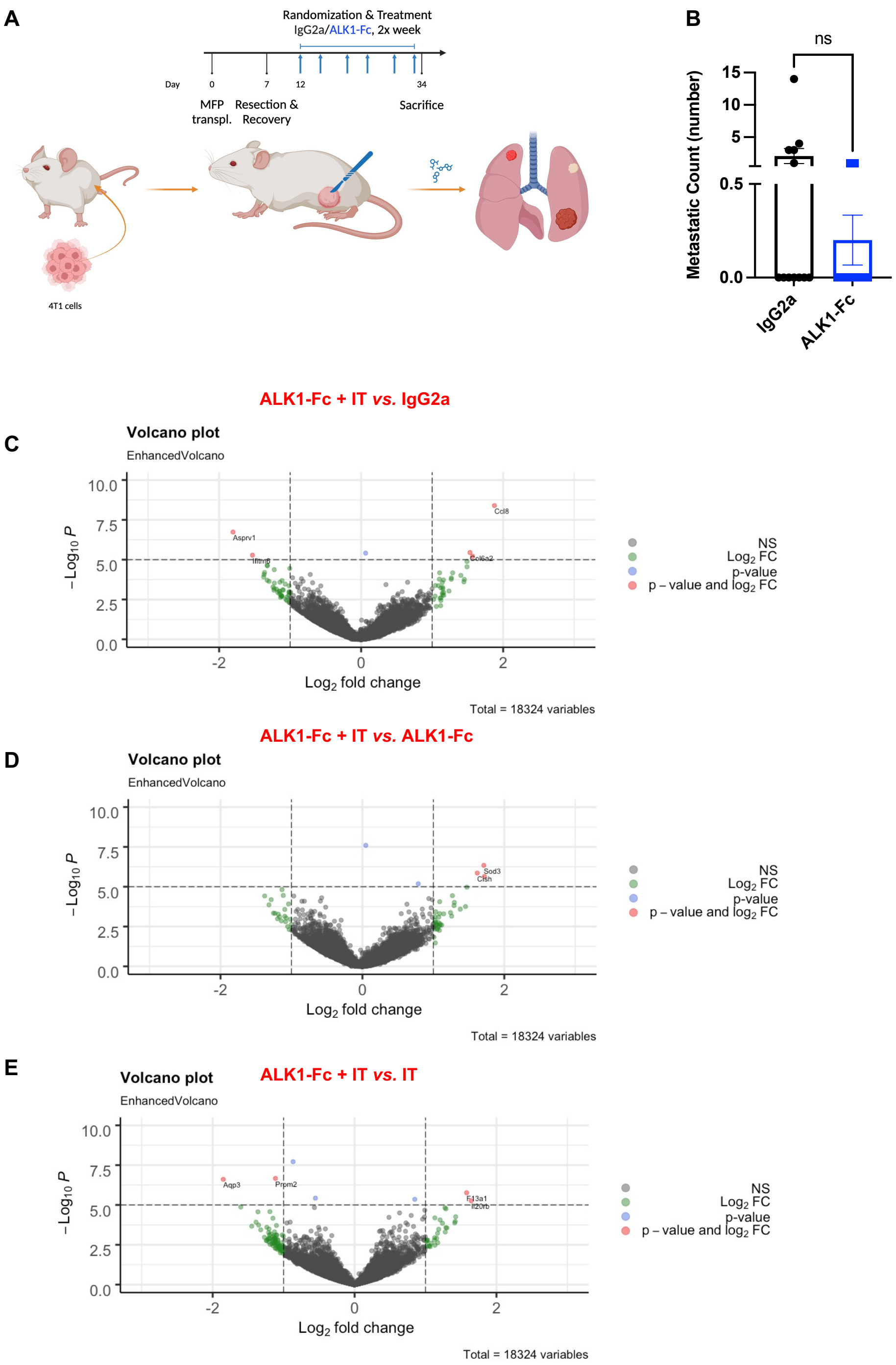

#### Figure S4.

(A): Experimental design of the adjuvant trial based on the orthotopic transplantation of $1x10^5$ 4T1 cells in syngeneic BALB/c hosts. Tumors were resected when reaching 4.5 mm in diameter. Mice were let to rest for 5 days before randomization. The groups received either ALK1-Fc or control IgG2a, twice weekly for 3 weeks. (B): Quantification of macro-metastases at the experimental endpoint, ALK1-Fc *vs* IgG2a. Data displayed as mean with SEM. p-value: Mann-Whitney U-test. (C-E): volcano plots from the bulk RNA-seq of the different treated cohort in the 4T1 adjuvant trial: ALK1-Fc + immunotherapy *vs* IgG2a (C), ALK1-Fc + immunotherapy *vs* ALK1-Fc (D), and ALK1-Fc + immunotherapy *vs* immunotherapy (E).

### Figure S5. Related to Figure 5

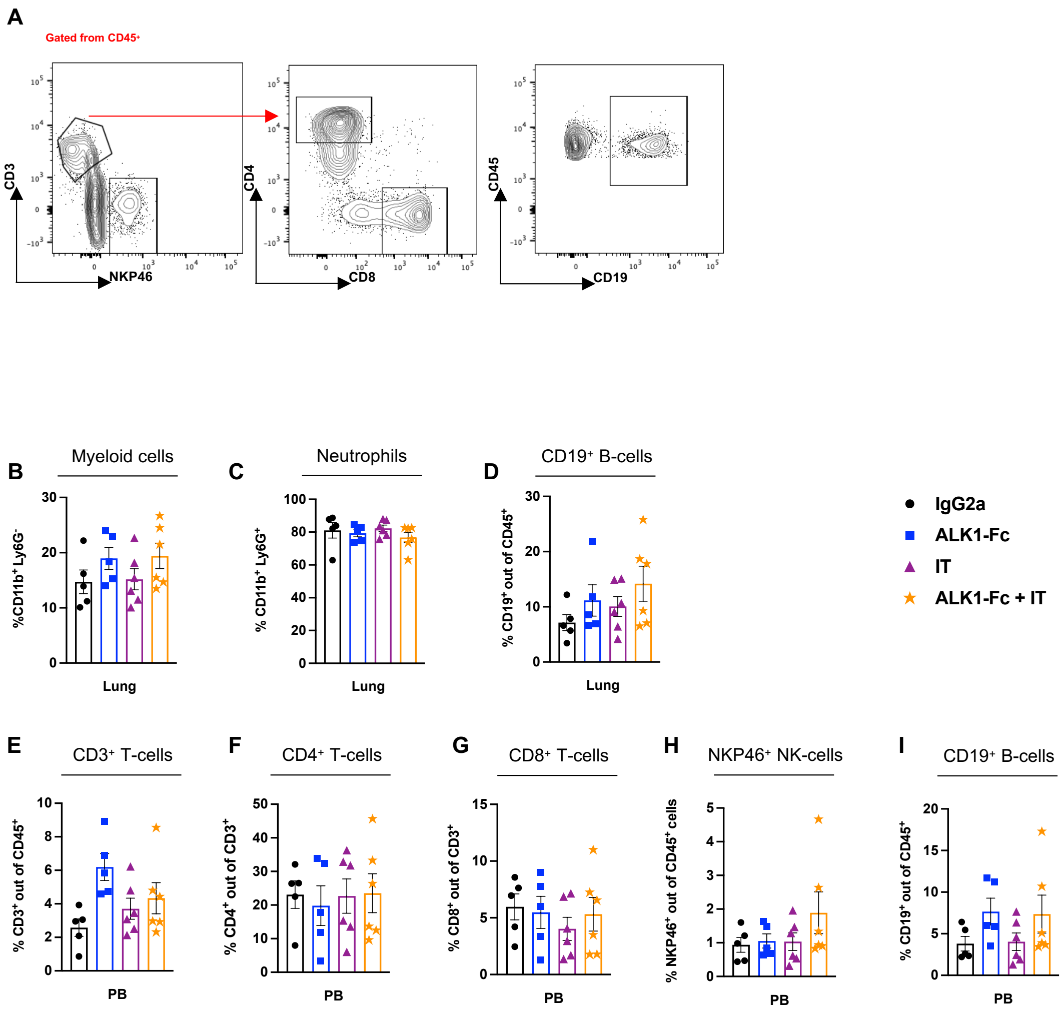

Figure S5.

(A): Gating strategy for lymphoid cells in lung tissue in the 4T1 adjuvant setup. (B-D): Relative frequency of myeloid cells (B), neutrophils (C), and B-cells (D) from lung tissue. Data displayed as mean with SEM. p-value: unpaired, two-tailed t-test. (E-I): Relative frequency of lymphoid cells in peripheral blood: $CD3^+$ T-cells (E), $CD4^+$ T-helper cells (F), $CD8^+$ cytotoxic T lymphocytes (G), $NKP46^+$ NK cells (H), and $CD19^+$ B-cells (I). Data displayed as mean with SEM. p-value: unpaired, two-tailed t-test.

### Figure S6. Related to Figure 7

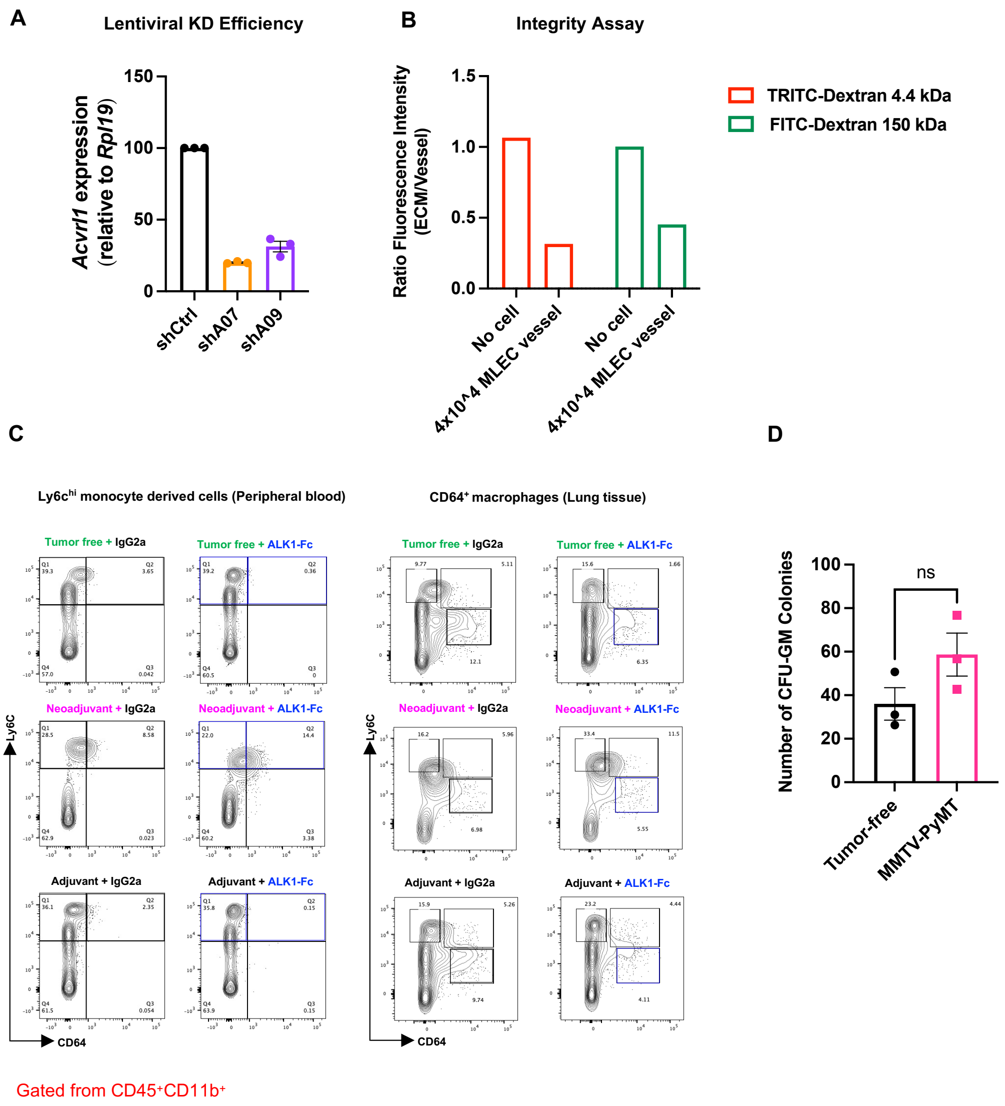

Figure S6.

(A): Lentiviral *Acvrl1* knockdown (KD) efficiency in mLEC cells. (B): Fluorescent labelled dextran was used to determine the vascular permeability of the mLEC endothelial tube in the 3D organ-on-a-chip setup. (C): Purity check of $Ly6C^{hi}$ monocyte-derived cells in peripheral blood (left) and $CD64^{+}$ macrophages in lung tissue (right) from the 4T1 short inhibition trial. (D): quantification of colony formation assay from $cKit^{+}$-enriched progenitor cells from the bone marrow of transgenic MMTV-PyMT tumor model (magenta) and littermate control (gray) mice. Data displayed as mean with SEM. p-value: paired, two-tailed t-test (ns: not significant).
